# Intra-dimer cooperativity between the active site cysteines during the oxidation of peroxiredoxin 2

**DOI:** 10.1101/2020.05.11.087908

**Authors:** Alexander V Peskin, Flávia C Meotti, Luiz F de Souza, Robert F Anderson, Christine C Winterbourn, Armindo Salvador

## Abstract

Peroxiredoxin 2 (Prdx2) and other typical 2-Cys Prdxs function as homodimers in which hydrogen peroxide oxidizes each active site cysteine to a sulfenic acid which then condenses with the resolving cysteine on the alternate chain. Previous kinetic studies have considered both sites as equally reactive. Here we have studied Prdx2 using a combination of non-reducing SDS-PAGE to separate reduced monomers and dimers with one and two disulfide bonds, and stopped flow analysis of tryptophan fluorescence, to investigate whether there is cooperativity between the sites. We have observed positive cooperativity when H_2_O_2_ is added as a bolus and oxidation of the second site occurs while the first site is present as a sulfenic acid. Modelling of this reaction showed that the second site reacts 2.2 ± 0.1 times faster. In contrast, when H_2_O_2_ was generated slowly and the first active site condensed to a disulfide before the second site reacted, no cooperativity was evident. Conversion of the sulfenic acid to the disulfide showed negative cooperativity, with modelling of the exponential rise in tryptophan fluorescence yielding a rate constant of 0.75 ± 0.08 s^-1^ when the alternate active site was present as a sulfenic acid and 2.29 ± 0.08-fold lower when it was a disulfide. No difference in the rate of hyperoxidation at the two sites was detected. Our findings imply that oxidation of one active site affects the conformation of the second site and influences which intermediate forms of the protein are favored under different cellular conditions.

## 1. Introduction

Peroxiredoxins (Prdxs) are a ubiquitous family of thiol proteins that react rapidly with hydrogen peroxide (H_2_O_2_). They are redox regulatory proteins that can function as antioxidants and as sensors in redox regulated signalling pathways. Prdx2 is a member of the typical 2-Cys (Prx1) subgroup, in which the functional unit is a non-covalent homodimer containing two active sites. The dimers reversibly associate to form decamers (1,2). The active sites each contain a highly reactive peroxidatic Cys residue (C_P_). These react independently with H_2_O_2_ to form a sulfenic acid. The sulfenic acid then condenses with the resolving Cys (C_R_) of the opposing subunit within the dimer to form a disulfide. This reaction occurs in competition with the reaction of the sulfenic acid with a further H_2_O_2_ to give the sulfinic acid. Disulfide formation is readily reversible, with the thioredoxin system being the most favored reductant. Hyperoxidation to the sulfinic acid is slowly reversed in an ATP-dependent process catalysed by sulfiredoxin.

Kinetic analyses to date have treated both active sites as equally reactive, such that modification of one site does not influence the reactivity of the other. In the current investigation, we have tested this assumption. A full scheme showing the individual reaction steps of the oxidation mechanism is shown in Fig 1, along with the designation of the symbols used to identify the different Prdx oxidation states and rate constants. We have reacted reduced Prdx2 with H_2_O_2_ and followed the formation of the one and two disulfide forms of the dimer by non-reducing SDS-PAGE as in (3–6), as well as disulfide and sulfinic acid formation by stopped flow monitoring of tryptophan fluorescence as in (7–9). These data have been fitted to kinetic models that account for any influence of the redox state at one site in a Prdx2 dimer on the oxidation, condensation, sulfinylation and reduction rates at the other site. This allowed us to determine the attending rate constants and characterize various modes of cooperativity between the two sites. With this approach we have been able to show that when substoichiometric concentrations of H_2_O_2_ are rapidly added as a bolus, oxidation of the second active site occurs while the first site is present as a sulfenic acid, and the second site reacts faster than the first. In contrast, with slow continuous generation of H_2_O_2_, the first active site condenses to a disulfide before the second site reacts and no cooperativity was evident. Analysis of changes in Trp fluorescence due to resolution of the sulfenic acid revealed negative cooperativity for disulfide formation and no cooperativity for sulfinic acid formation.

**Figure 1.**
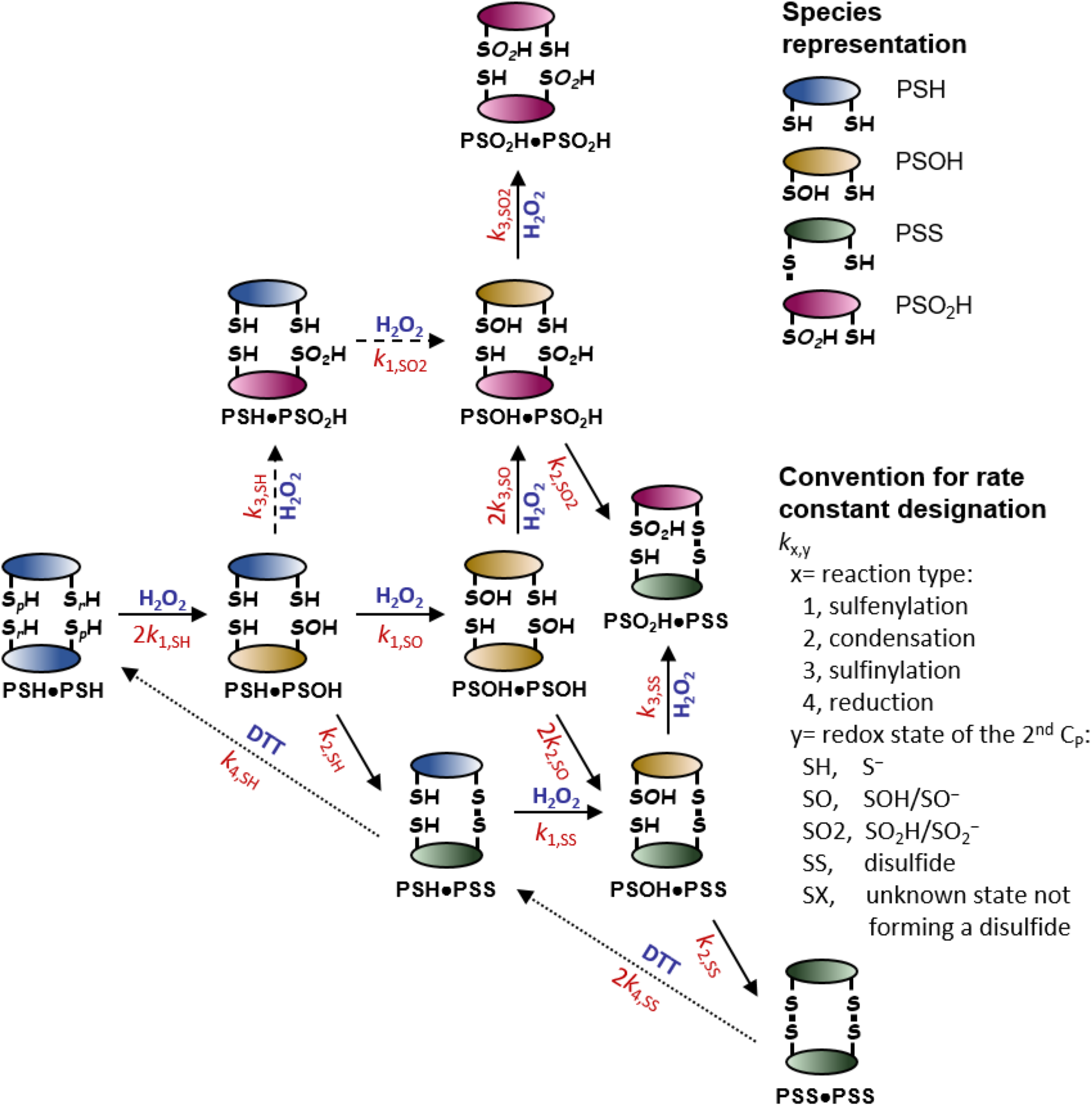
Single step scheme for Prdx2 oxidation within the functional dimer. Horizontal, solid diagonal and vertical arrows represent sulfenylation, condensation and sulfinylation steps, respectively. The dashed steps represent minor oxidation pathways. Rate constants are represented in red. The dotted steps represent reduction by DTT. Sulfenic and sulfinic acids are represented protonated for typographic convenience. None of the models used below makes a distinction between the properties of protonated and unprotonated forms.

## 2. Materials and Methods

### 2.1 Materials

Human recombinant WT and C172S Prdx2 (untagged) were prepared in the absence of DTT as described (10) and kindly provided by Dr Paul Pace. Reduced Prdx2 was prepared by incubation with 10 mM DTT for 1 h in pH 7.4 phosphate buffer, 0.14 M NaCl (PBS) followed by removal of the reductant using a Micro Bio-Spin 6 column (Bio-Rad). Columns were washed and pre-equilibrated with 20 mM phosphate buffer pH 7.4 containing 0.1 mM diethylenetriamine-penta-acetic acid (DTPA) purged with argon. Protein concentration was determined by measuring A_280_ using a NanoDrop spectrophotometer (Biolab Nanodrop Technologies, Wilmington, DE) and ε_280_=21430 cm^-1^M^-1^, or with a DirectDetect spectrometer (Merck). Hydrogen peroxide (Fluka) was standardized by measuring A_240_ (ε = 43.6 M^-1^cm^-1^).

### 2.2 Prdx2 treatment and SDS-PAGE analysis

Several methods were employed for bolus addition of H_2_O_2_ to Prdx2. Standard rapid mixing was performed in Eppendorf tubes in total volumes varied from 10 μl to 40 μl. Reactions were performed I 20 mM phosphate buffer pH 7.4 containing 0.1 mM DTPA. Either protein or H_2_O_2_ was placed as a drop on the wall of the tube and then sharply blasted with the rest of the mixture placed at the bottom using a vortex mixer. The reaction was stopped by adding 10 mM *N*-ethylmaleimide (NEM) for SDS-PAGE. Rapid mixing was also performed with the stopped flow system as used for measuring tryptophan fluorescence. The reaction mixture flowing out from the stopped flow line was collected in a tube containing 10 mM NEM. The 2 mL collected was concentrated in a speed vacuum to 100-200 μL.

Mixing was avoided entirely by exposing 10 μM Prdx2 to substoichiometric H_2_O_2_ generated in situ using pulse radiolysis. The University of Auckland’s 4 MeV linear accelerator was used to deliver a range of radiation doses (3-50 Gy), in 2 μs to 100 μL samples in glass tubes. Radiation doses per pulse over a range of machine settings, were calibrated using FRICKE dosimetry assuming a G-value for the production of Fe^3+^ ions of 1.606 μM.Gy^-1^. Dose-dependent superoxide concentrations were produced in aerated solutions containing sodium formate (50 mM), DTPA (0.1 mM) and phosphate buffer (20 mM) at pH 7. Concentrations of superoxide were calculated to be 0.62 μM.Gy^-1^ and found to be in agreement with superoxide concentrations scavenged by added cytochrome c (50 μM), utilizing Δε_550nm_ = 27,600 M^-1^ cm^-1^. Hydrogen peroxide concentrations were produced by the pulse radiolysis of the above solution with added Cu,Zn-superoxide dismutase (5 μM) and calculated to be half the concentration of superoxide (0.62 μM.Gy^-1^/2) plus the molecular yield of H_2_O_2_ (0.07 μM.Gy^-1^) produced at each dose setting. Hydrogen peroxide concentrations produced per pulse were checked using the Amplex Red assay. Pre-existing H_2_O_2_ in buffers was removed by incubating with 2 mg/ml catalase in a dialysis bag.

To expose Prdx2 to continuous H_2_O_2_ generation the protein (20 μM) was incubated with 5 mM glucose plus glucose oxidase (Sigma). Glucose oxidase was diluted to produce approximately 0.5 μM H_2_O_2_/min.

Prdx2 species were separated by non-reducing SDS-PAGE on 10% gels (3,5), visualized by Coomassie staining, and documented with an Alliance Q9 Advanced Chemiluminescence Imager (Bio-Rad). The relative intensities of the Prdx2 bands were quantified by densitometry using Quantity One software (Bio-Rad).

### 2.3 Stopped flow analysis of tryptophan fluorescence

For kinetic measurements based on intrinsic fluorescence changes (7–9), Prdx2 was reduced with a five-fold excess of DTT per cysteine the day before experiment. The remaining DTT was removed by filtration (Amicon Ultra 10 kDa, Merk Millipore, Darmstadt, Germany). The protein solutions were stored in an oxygen-free atmosphere. Oxidation of pre-reduced Prdx2 (~2.5 μmol SH×μmol protein^-1^) or C172S Prdx2 (~1.6 μmol SH×μmol protein^-1^) by increasing concentrations of H_2_O_2_ was followed by mixing in a stopped-flow instrument (Applied Photophysics *SX20MV, Leatherhead, United Kingdom*), excitation λ_280nm_, emission above λ_320nm_. The reactions were performed at 25 °C in 5 mM sodium phosphate buffer pH 7.4 containing 100 μM DTPA. Buffer solution was previously treated with 10 μg/mL catalase to remove any trace of H_2_O_2_ (5). Reactions were ran at sub-stoichiometric or supra-stoichiometric concentrations of H_2_O_2_.

### 2.4 Data analysis

Details of the modelling and data analysis are presented in the Supplementary Material (SM). Briefly, the cooperativity parameters *R*_1,SS_ and *R*_1,SO_ were determined by fitting the numerical solutions of models of the reaction schemes in Figs 4A and 4B, respectively, to the gel densitometry data (SM2.1.1 and 2.1.2). Due to the presence of significant initial fractions of one-disulfide (1DS) and anomalous dimers (see below) in some Prdx2 solutions slightly more complex models had to be used to determine *R*_1,SO_ that accounted for the reactivity of these dimers. To ensure that each dataset was fitted by the best suitable statistical model we systematically tested a family of 24 models embodying distinct assumptions about the reactivity and initial distribution of anomalous dimers between the monomer and 1DS bands and distinct sets of adjustable parameters (details in SM2.1.2). For each data set we selected among the statistical models that yielded best fit estimates significantly different from 0 for all the adjustable parameters the one with the lowest value of the Akaike Information Criterion (11) with small-sample correction.

In order to fit the Trp fluorescence recovery time series we first selected data in a time range 0.2 s after the fluorescence minimum to 75% of the time of the fluorescence maximum. This time window avoids both the initial concave-up phase of the fluorescence recovery and the late phase where bleaching becomes relevant. We then applied an iterative approach fitting the selected data with linear combinations of an increasing number of exponentials. Linear combinations of *n* exponentials were judged the best statistical models of the data whenever the following four criteria were cumulatively satisfied: (i) the fits yielded plausible estimates and tight confidence intervals for all adjustable parameters; (ii) systematic bias in the fit residuals were eliminated or attenuated relative to *n*-1-exponential fits; (iii) the Akaike Information Criterion had a lower value than those for *n*-1- and *n*+1-exponential fits; (iv) the parameter estimates were consistent across replicates. Further details of the procedures above are provided in SM2.2.1.

## 3. Results

### 3.1 Positive cooperativity in the reaction of reduced Prdx2 with bolus H_2_O_2_

To follow Prdx2 oxidation by H_2_O_2_, the H_2_O_2_ was either generated slowly with glucose/glucose oxidase or added as a bolus, and the reaction was followed by blocking reduced thiols and separating the reaction mixture using non-reducing SDS-PAGE. As shown in Fig 2, the Prdx2 was initially almost completely reduced and ran as a monomer, and H_2_O_2_ treatment resulted in the formation of two dimer bands. The upper band has been shown to correspond to dimer with one disulfide bond, and the lower band to have two disulfide bonds (5,12). When H_2_O_2_ was generated continuously (~1.7 nM/s), Prdx2 oxidation occurred progressively over time (Fig 2A). It is apparent from the gel that the 1DS band gradually built up and then the two disulfide (2DS) band accumulated. A similar pattern of oxidation to Fig 2A was also seen when reduced Prdx2 was stored over several days and gradually oxidized by adventitious peroxides that formed in the solution (not shown).

**Figure 2.**
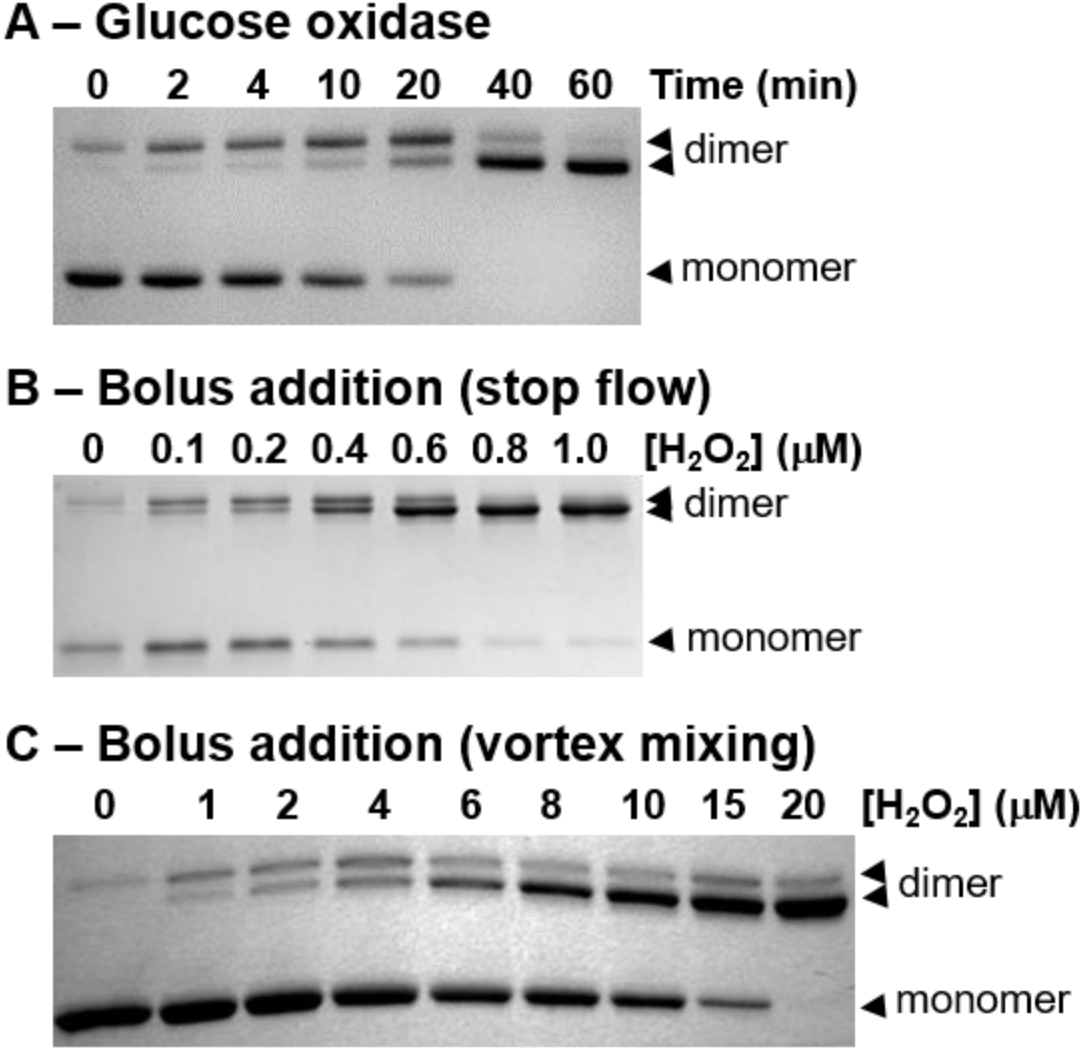
Formation of dimers containing one and two disulfides on the oxidation of Prdx2 by H_2_O_2_. (**A**) Representative gel showing oxidation profile over time when Prdx2 (20 μM) was incubated with glucose/glucose oxidase to generate H_2_O_2_ at ~1.7 nM/s, with reaction stopped by adding NEM at stated times. (**B**) Representative gel showing oxidation profile after stopped flow mixing of 1.0 μM Prdx2 with indicated concentrations of H_2_O_2_ and collecting into NEM. (**C**) Representative gel for treatment of 20 μM reduced Prdx2 with indicated concentrations of H_2_O_2_ using vortex mixing. NEM was added after 5 min. Samples were subjected to non-reducing SDS/PAGE. Coomassie stained gels are shown.

Treatment of reduced Prdx2 with bolus additions of H_2_O_2_ up to a 1:1 mole ratio gave a more complex picture. With conventional rapid mixing of the two reactants by vortexing, the two-disulfide forms started to accumulate even at the lower concentrations and the one-disulfide form never built up (Fig 2B). However, because the reaction is so fast, preferential oxidation of the second site could be the result of limited diffusion during mixing. To test this, we used the stop flow mixing system, as described below and widely used to determine rate constants for the reaction, and analysed the Prdx2 redox state in the flow through. The gel suggests that there was preferential oxidation, but not as much (Fig 2C). We also eliminated the mixing factor by using pulse radiolysis to generate various amounts of H_2_O_2_ *in situ*. This gave a similar gel pattern to that seen in Fig 2C.

To analyse these data, densitometry measurements of the percentages of each component were made for multiple experiments and the data plotted as a function of the fraction (f) of initially reduced sites that were oxidized (Fig 3). Simple probabilistic considerations show that if the two sites are equivalent and independently oxidized, the fractions of dimers with zero, one and two disulfides would fit the curves shown as dashed lines. The time course data for continuous generation fit well to this random mechanism (Fig 3, top). There is a poorer fit for the data obtained with stopped flow mixing (bottom) and there was greater divergence for the vortex-mixed samples (not shown).

**Figure 3.**
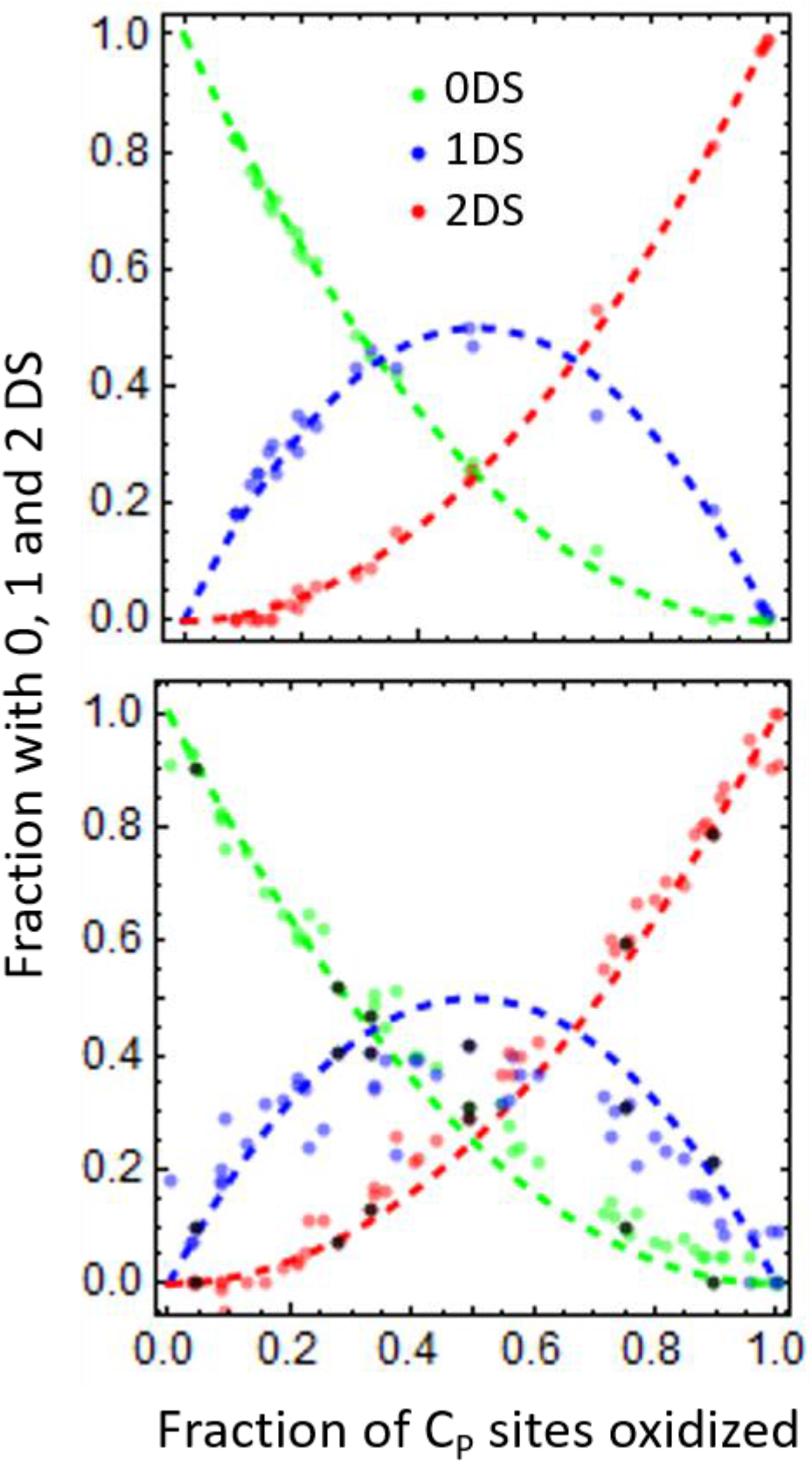
Kinetic analysis of gel data to assess cooperativity in the oxidation of reduced Prdx2. Analysis of densitometry data. Upper panel: Reduced Prdx2 (~10 μM) was oxidized by slow (***v***_sup_= 0.87 nM/s, 1.7 nM/s and 3.6 nM/s) continuous generation of H_2_O_2_ as in Fig 2A (results from 3 independent experiments). Lower panel: Reduced Prdx2 (1.0-2.0 μM) was mixed with various substoichiometric amounts of H_2_O_2_ by stopped flow (results from 3 independent experiments) and pulse radiolysis (black dots). Proportions of reduced monomer (green symbols), 1DS dimers (blue symbols), and 2DS dimers (red symbols) are shown for each experimental result. The broken lines in the respective colors show the binomial distribution curves *f*_0_= (1-*f*)^2^, *f*_1_= 2 *f*(1-*f*) and *f*_2_=*f*^2^, where *f* is the fraction of C_P_ sites that were oxidized and the subscripts correspond to the number of disulfides) expected if a thiolate in a 1DS dimer (top) or one-sulfenate (bottom) is oxidized at the same rate as that in the two-thiolate dimer. Only the slow generation results fit well with this expectation. The bolus addition results yield a higher proportion of 2DS vs. 1DS dimers. See text for further explanation.

More-sophisticated simulations to quantify the extent of cooperativity drew on the dimer-based reaction scheme for Prxd2 oxidation from Benfeitas et al. (13) (Fig 1). Here, the oxidation of C_P_ by H_2_O_2_ and the condensation with C_R_ are considered independently for each active site. Thus there are two possibilities for C_P_ oxidation at the second site, depending on whether the first site is a sulfenic acid or whether it has already condensed to the disulfide. The two experimental systems examined represent extreme situations, which simplifies analysis. With continuous generation of H_2_O_2_ at <5 nM/s and ≥10 μM Prdx2, the C_P_ are sulfenylated with a pseudo-first-order rate constant <0.002 s^-1^ (SM2.1.1). Therefore, the chances of H_2_O_2_ hitting the second site before the first sulfenic acid condensed should be low. The relevant reaction scheme for modelling these experiments can thus be simplified as indicated in Fig 4A. The symbols and rate constants are defined in Fig 1. In contrast, because the reaction of H_2_O_2_ with the C_P_ thiols of Prdx2 is very fast (k >10^7^ M^-1^s^-1^) and the rate of condensation of the sulfenic acid and hyperoxidation relatively slow — rate constants of 0.2-1 s^-1^ and <6000 M^-1^s^-1^ respectively have been reported (5,9,14,15) — all the H_2_O_2_ added as a bolus should oxidize the second site while the first site is a sulfenic acid. The relevant reaction scheme can thus be simplified as indicated in Fig 4B. The methodology used for analysis of these two situations is given in the Data Analysis section, with more detail in SM2.1.

The main parameters relating to cooperativity are the relative sulfenylation rates for the second site compared with the first site, *R*_1,SS_= *k*_1,SS_/*k*_1,SH_ for the disulfide and *R*_1,SO_= *k*_1,SO_/*k*_1,SH_ for the sulfenic acid. The data for each experiment used to create Fig 3 were modelled on this basis to give a best fit for *R*_1,SS_ or *R*_1,SO_. An example of the fits for slow generation is shown in Fig 4C. Best-fit values for *R*_1,SS_ obtained from 3 independent slow generation or spontaneous oxidation experiments were all close to 1 (as expected from the good fit to the random distribution curves in Fig 3), yielding a variance-weighted mean 1.25±0.07. In contrast, best-fit values for *R*_1,SO_ obtained from 6 independent stopped flow mixing experiments (an example shown in Fig 4D) were in the range 2.0+0.4 to 5.3+0.8 giving a variance-weighted mean for *R*_1,SO_ of 2.3±0.2 (95% confidence interval [2.0, 2.7]). The pulse radiolysis data gave a *R*_1,SO_ value of 2.25+0.05, i.e. within the stopped flow range (variance-weighted mean of all these experiments: 2.2±0.1). These results are therefore indicative of the second site reacting about twice as fast as the first.

**Figure 4.**
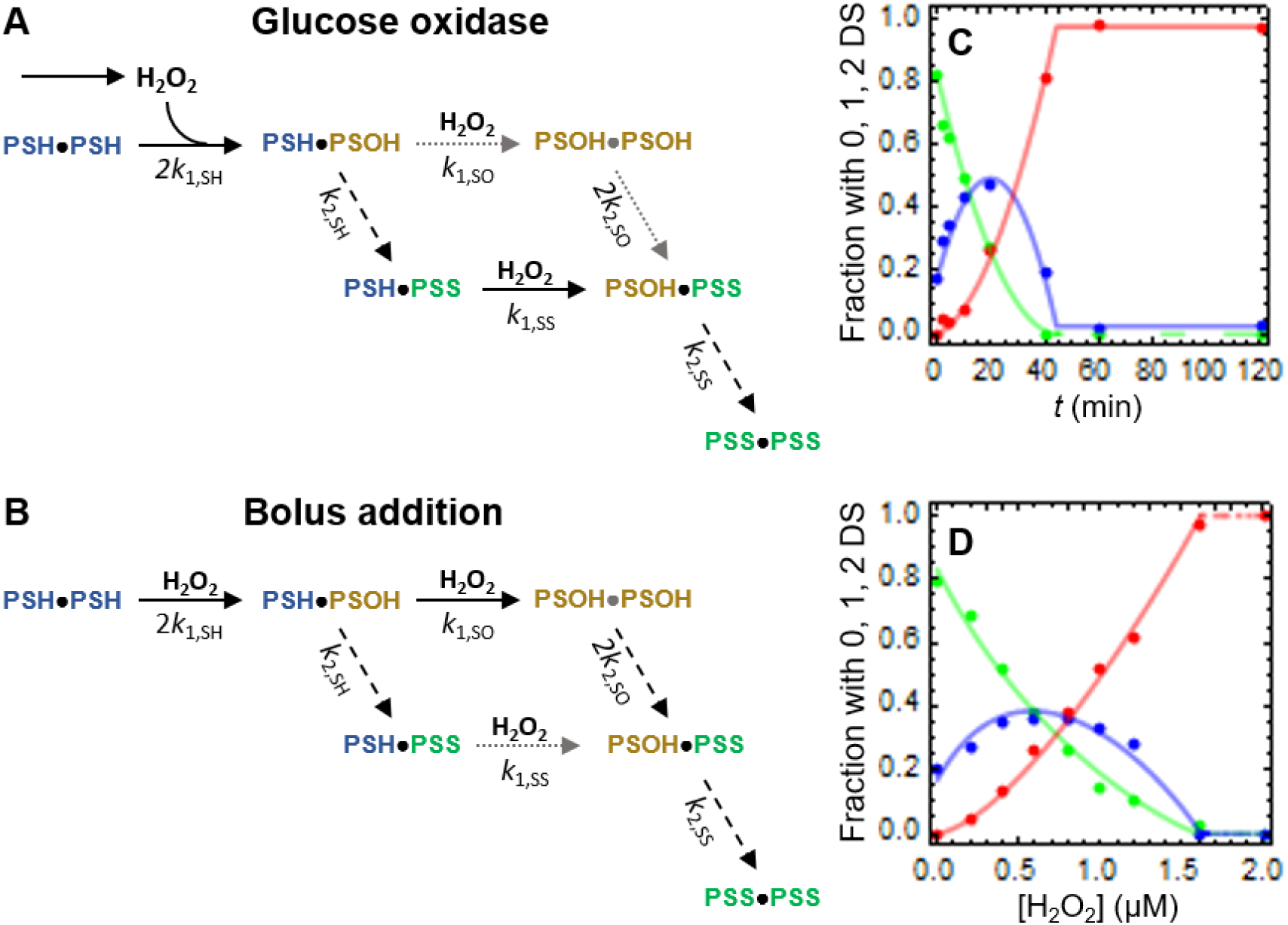
Underlying model for the reaction of sub-to stoichiometric H_2_O_2_ with the two active sites and representative fits. (**A**) When H_2_O_2_ is supplied at nM/s rates to μM Prdx2, two-thiolate Prdx2 dimers (PSH·PSH, migrating on the gel as monomers) are converted to one-sulfenate dimers (PSH·PSOH). The first sulfenic acid has time to condense before the second C_P_ is oxidised. Thus, the pathway indicated by gray dotted arrows does not significantly contribute for disulfide formation, and rate constants *k*_1,SO_, *k*_2,SO_ virtually do not influence the final outcome. Neither do the rate constants *k*2,SH and *k*2,SS (dashed arrows), because the formation of the disulfides is rate-limited by the sulfenylation steps (solid black arrows). The distribution of monomers, one- and two-disulfide dimers at each point in time is thus solely determined by the overall extent of Prx2 oxidation and by the relative oxidation rate (*R*_1,SS_= *k*_1,SS_/*k*_1,SH_) of a thiolate in a 1DS dimer *vs*. a thiolate in a two-thiolate dimer. The value of *k*_1,SH_ has no influence, and only the two reactions indicated by solid black arrows need to be explicitly considered in modelling this distribution. (**B**) When μM H_2_O_2_ boluses are added, PSH·PSH is converted to one- (PSH·PSOH) and then two-sulfenate (PSOH·PSOH) dimers (solid black arrows) in milliseconds, before the first sulfenate condenses. Thus, the reaction indicated by a gray dotted arrow contributes little to the formation of the 2DS dimers (PSS·PSS), and the respective rate constant (*k*_1,SS_) virtually does not influence the final outcome. Neither do the rate constants *k*_2,SO_, *k*_2,SS_ and *k*_2,SH_ (dashed arrows), because the two-sulfenate dimers are committed to form 2DS dimers, and the one-sulfenate dimers left after the H_2_O_2_ vanished are committed to form 1DS dimers (PSH·PSS). The distribution of monomers, one- and two-disulfide dimers is thus solely determined by the H_2_O_2_:Prdx2 stoichiometry and by the relative oxidation rate (*R*_1,SO_= *k*_1,SO_/*k*_1,SH_) of a C_P_ in a one-sulfenate dimer *vs*. a C_P_ in a two-thiolate dimer. The value of *k*_1,SH_ has no influence, and only the two reactions indicated by solid black arrows need to be explicitly considered in modelling this distribution. (**C**, **D**) Representative fits (lines) of the models described in (A) and (B) to experimental data (dots) from slow H_2_O_2_ generation and bolus addition through a stopped flow apparatus, respectively. Proportions of reduced monomer (green), 1DS dimers (blue), and 2DS dimers (red) are shown.

Analysis of the data from 9 experiments with vortex mixing gave a variance-weighted mean value for *R*_1,SO_ of 4.9±0.1 (95% confidence interval [4.7, 5.1]), which is significantly higher than for stopped flow mixing. This suggests that part of the apparent cooperativity is due to limited diffusion with this type of mixing. Consistent with this hypothesis, the values of *R*_1,SO_ obtained with various volumes and concentrations of reactants showed a strong positive correlation with the concentration of Prdx2 used (σ=0.83, p<5×10^-6^, not shown).

It should be noted that a small proportion of the reduced starting material (typically around 10%) was frequently present in the upper Prdx2 (1DS) dimer band. This could represent dimer with one reduced active site, arising from incomplete reduction or a small amount of oxidation occurring during the removal of DTT. Also, in some experiments a 1DS dimer band remained after stoichiometric H_2_O_2_ had been added. This band was insensitive to oxidation and therefore could not represent dimer with one reduced active site. Although in theory it could represent dimer with one hyperoxidized active site, hyperoxidation was not detected in the starting material by LC/MS analysis and only became evident after treatment with higher H_2_O_2_ concentrations. We presume that the band represents a form of dimer where one of the sites (designated X) was unable to form a C_P_-C_R_ disulfide (henceforth called “anomalous dimers”) but have not pursued its identity further. The presence of this form could be accounted for by using a slightly more complex model (SM2.1.2 and Fig S1), but this yielded best-fit *R*_1,SO_ estimates in the same range as the other experiments.

### Negative cooperativity for disulfide formation

Prdxs contain tryptophan residues that change in fluorescence intensity during oxidation, thus enabling the oxidation process to be monitored by stopped flow (7–9). A typical trace shows a rapid decrease in fluorescence associated with the initial oxidation step followed by a slower recovery phase (Fig 5A). In previous studies these have been analyzed as single exponentials to obtain rate constants for the two steps. In the current study we have carried out more rigorous data analysis to test if the two sites can be distinguished kinetically. To do this requires consideration not only of the rate constants for the different reactions, but also the fluorescence quantum yields for each species in the reaction sequence, thus making analysis more complex.

**Figure 5.**
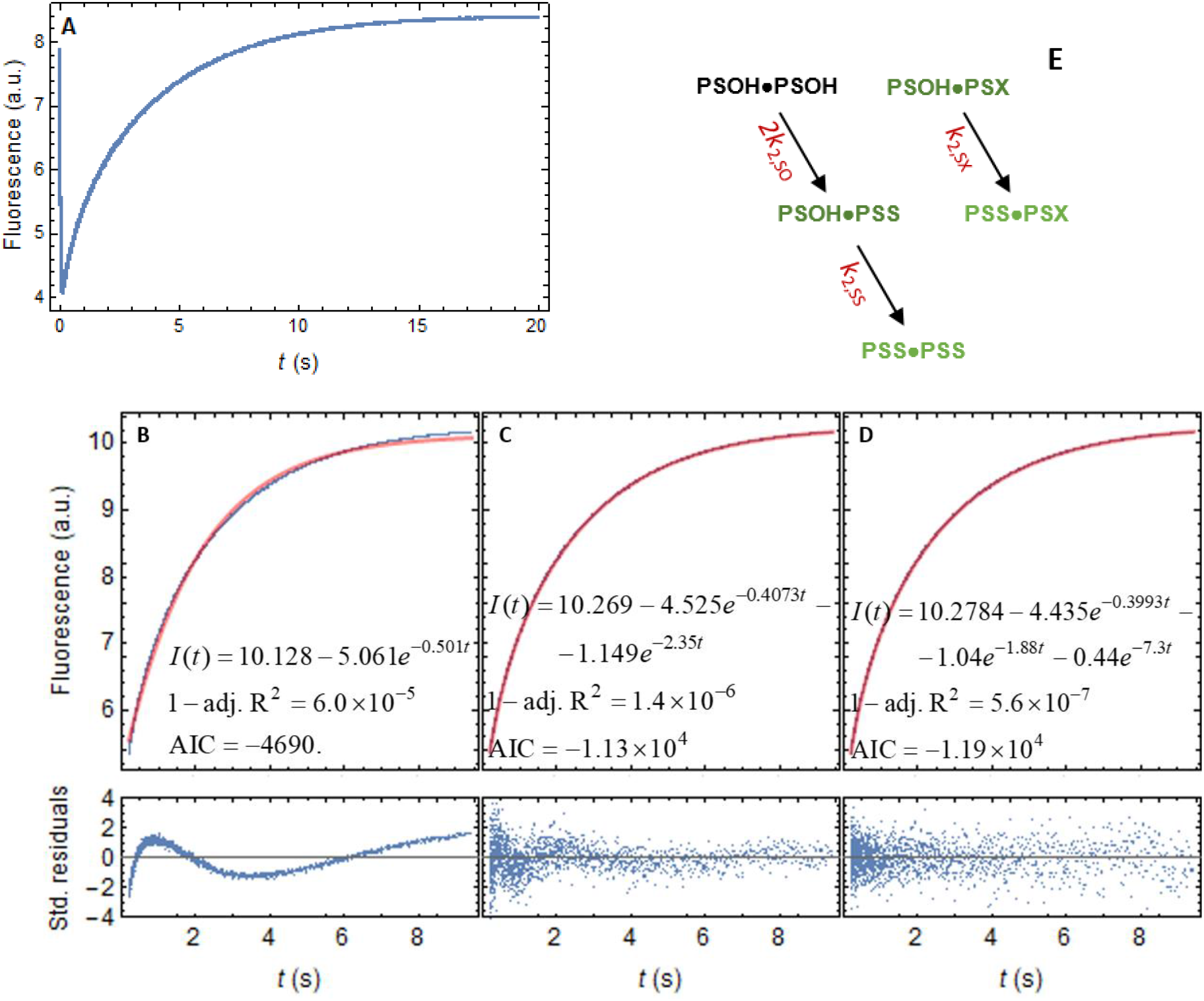
Kinetic analysis of Trp fluorescence recovery after Prdx2 oxidation by stoichiometric H_2_O_2_. (**A**) Full fluorescence time course for treatment of 0.5 μM Prdx2 with stoichiometric H_2_O_2_, generated as described in Methods section. (**B**) Mono-, (**C**) bi- and (**D**) tri-exponential variance weighted fits to a typical Trp fluorescence recovery time course (blue) after treatment of 1.3 μM Prdx2 monomers with 1.6 μM H_2_O_2_. Red, best-fit curves. A systematic pattern of deviations of the fluorescence from the mono- and bi-exponential best-fit fits curves (B, C, bottom panels) is consistent among replicates, revealing that fluorescence recovery is more complex. A tri-exponential fit eliminates the systematic deviation (D, bottom panel). The tri-exponential model has significantly higher likelihood (p<10^-120^) than the bi-exponential model. (**E**) Reaction scheme to simulate the fluorescence recovery. At the μM H_2_O_2_ concentrations used in the Trp quenching experiments sulfenic acid formation is much faster than condensation. For analyzing the kinetics of condensation it is thus a good modelling approximation to consider that all the sulfenic acids are instantaneously formed. The condensation of sulfenic acids from a small fraction of anomalous dimers (PSOH·PSX) is hypothesized to contribute to fluorescence recovery. Darker symbols indicate lower fluorescence quantum yields (PSOH·PSOH < PSOH·PSS ≈ PSOH·PSX < PSS·PSS ≈ PSS·PSX).

We first tried to examine the rapid phase. Unfortunately, even the simplest models for formation of the sulfenic acids in a dimer imply several intermediate species whose fluorescence quantum yields are unknown and have to be treated as adjustable parameters in the analyses. These uncertainties combined with the inherent noisiness of the data, meant that cooperative and non-cooperative models could be fitted to the data and robust conclusions about cooperativity could not be drawn.

We then analysed the Trp fluorescence recovery after Prdx2 oxidation, which can be attributed to condensation of the sulfenic acid to form the disulfide. Mono-exponential fits with stoichiometric H_2_O_2_ left a well-defined pattern of residuals (Fig 5B) that was consistent over replicates and experiments. A detailed analysis through the approach explained in SM2.2.1 revealed a tri-exponential recovery, with all the pre-exponential coefficients negative (Fig 5B-D, SM2.2.2). The characteristic constants and scaled pre-exponential coefficients obtained are shown in Table 1, experiment a. A tri-exponential fluorescence recovery when only two sulfenic species -- PSOH·PSOH and PSOH·PSS – were expected was surprising. However, it can be explained by the oxidation and subsequent condensation of the active site remaining in the “anomalous dimers” that were detected in the gel analysis of the flow through from the stopped flow apparatus. Indeed the model shown in Fig 5E fitted the data and provided a mechanistic interpretation for all the characteristic constants and scaled pre-exponential coefficients (details of analysis in SM2.2.2). In particular, this modelling supports the attribution of the rate constants to the condensation of the three species, PSOH·PSOH, PSS·PSOH and PSX·PSOH (rate constants *k*_2,SO_, *k*_2,SS_ and *k*_2,SX_ in Table 2, row a). An independent data set obtained with a different Prdx2 preparation yielded similar results (Tables 1,2, experiment b, see SM2.2.4 for details of analysis). In both cases the condensation rate constant for each sulfenic acid in a PSOH·PSOH dimer is a little over twice that for PSOH·PSS (*R*_2,SS_ = 0.42-0.48), revealing a negative cooperativity in condensation.

**Table 1:** Best-fit parameter estimates for the tri-exponential fits to fluorescence recovery time courses. Characteristic constants (*k*), scaled pre-exponential coefficients (*w*) and slopes of the characteristic constants with respect to H_2_O_2_ concentration (*sl*). Means ± standard errors from *n* replicates. Note that experiment *b* was done with a different Prdx2 preparation from *a* and gives slightly lower values for the characteristic constants. The characteristic constants correspond to *k*_2_, and the slopes in experiment c correspond to *k*_3_.

| Experiment | n | Parameters | Characteristic constants, weights and slopes |  |  |
| --- | --- | --- | --- | --- | --- |
|  |  |  | Exponential 1 | Exponential 2 | Exponential 3 |
| a. 1.3 $\mu\text{M}$ Prdx2 +<br>1.6 $\mu\text{M}$ $\text{H}_2\text{O}_2$, 26 $^\circ\text{C}$ | 7 | $k$ ($\text{s}^{-1}$)<br>$w$ | 0.396±0.004<br>0.74±0.01 | 1.88±0.03<br>0.178±0.003 | 7.9±0.7<br>0.09±0.01 |
| b. 0.5 $\mu\text{M}$ Prdx2 +<br>0.5 $\mu\text{M}$ $\text{H}_2\text{O}_2$, 24 $^\circ\text{C}$ | 4 | $k$ ($\text{s}^{-1}$)<br>$w$ | 0.2445±0.0004<br>0.791±0.001 | 1.07±0.03<br>0.139±0.001 | 3.28±0.04<br>0.0703±0.0006 |
| c. 0.5 $\mu\text{M}$ Prdx2 +<br>suprastoich. $\text{H}_2\text{O}_2$,<br>24 $^\circ\text{C}$ (extrapolated) | - | $k$ ($\text{s}^{-1}$)<br>$sl$ ($\text{M}^{-1}\text{s}^{-1}$) | 0.2435±0.0009<br>(3.33±0.03)$\times 10^3$ | 1.15±0.03<br>(8.±1.)$\times 10^3$ | 3.8±0.1<br>(18.±6)$\times 10^3$ |

Although there may be alternative explanations for the tri-exponential fluorescence recovery (see SM2.2.5), the following additional observations further support the cooperativity model. First, the results from the gel experiments with supra-stoichiometric H_2_O_2_ boluses described in the next section are consistent with this model and not with the alternative serial isomerizations model discussed in SM2.2.5. Second, all the three characteristic constants show a similar pH dependence (SM2.2.6 and Fig 6A) that is as expected for processes tightly coupled to condensation (15) and not for processes of a distinct nature. Indeed, the kinetic model for the pH dependence of the condensation rate constant presented in (15) fits the dependence of each of the three characteristic constants very well (Fig 6A). And the best fit estimates of pK_a_(C_P_-SOH) and pK_a_(C_R_-SH) are not significantly different among the characteristic constants (Table S2). This analysis also shows that the negative cooperativity in condensation is weakest at neutral pH and increases (*R*_2,SS_ decreases) towards more acidic and basic pH (Fig 6B).

**Figure 6.**
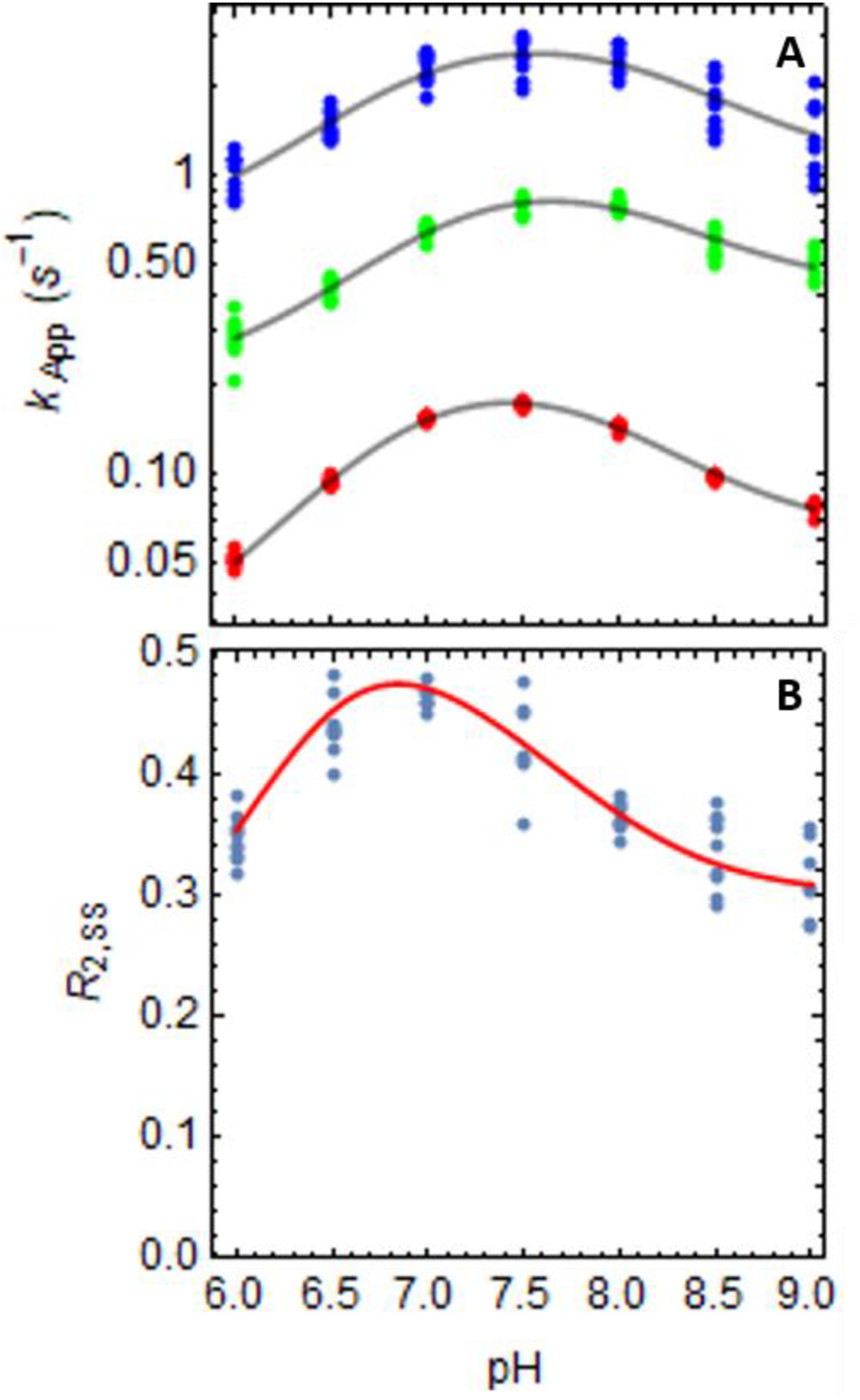
pH dependence of the characteristic constants for fluorescence recovery and of cooperativity in condensation. (a) pH dependence of the characteristic constants obtained from tri-exponential fits to the Trp fluorescence recovery after treatment of 1 μM Prdx2 with 1 μM or 2 μM H_2_O_2_ (representative experiment). The dots represent best-fit estimates of the characteristic constants for replicate injections from two experiments. The solid lines represent the best-fit curves for the pH-dependence model described in ref. (15). (B) pH dependence of the cooperativity in condensation. Dots, based on the *k*_App_ values in panel (A); line, calculated from the best-fit parameters for the pH-dependence model.

### No cooperativity for Prdx2 hyperoxidation

We also investigated site-to-site differences in susceptibility to hyperoxidation by exposing reduced Prdx2 to higher concentrations of H_2_O_2_. As hyperoxidation occurs in competition with disulfide formation (Fig 7A), variation in hyperoxidation between the sites could be due to either or both of these rates differing. Therefore, in order to sort out possible ambiguities we began by analyzing the hyperoxidation of a C_R_ mutant (C172S) Prdx2. This mutant cannot form interchain disulfides but the reduced form shows similar reactivity with H_2_O_2_ as the WT protein (6). Addition of H_2_O_2_ to reduced C172S Prdx2 causes a rapid drop in fluorescence attributable to oxidation of C_P_ to the sulfenic acid, followed by a gradual recovery that is attributable to sulfinylation (Fig 7B). Analysis of the data obtained with various supra-stoichiometric H_2_O_2_ concentrations [1-10 μM H_2_O_2_ over 0.5 μM Prdx2] using the approach described in SM2.2.1 reveals a bi-exponential recovery (Fig 7B), the slow component being due to bleaching (analysis in SM2.2.3). The single characteristic constant attributable to hyperoxidation shows a linear dependence on the H_2_O_2_ concentration with a slope (3.4±0.1)×10^3^ M^-1^V^-1^ (Fig 7C). Because under the conditions of the experiment all the peroxidatic cysteines are hyperoxidized [Figure 2c from (6)], the single characteristic constant implies that the formation of the first sulfinic acid does not influence the rate constant for the second sulfinylation. That is, *k*_3,SO_= *k*_3,SO2_.

**Figure 7.**
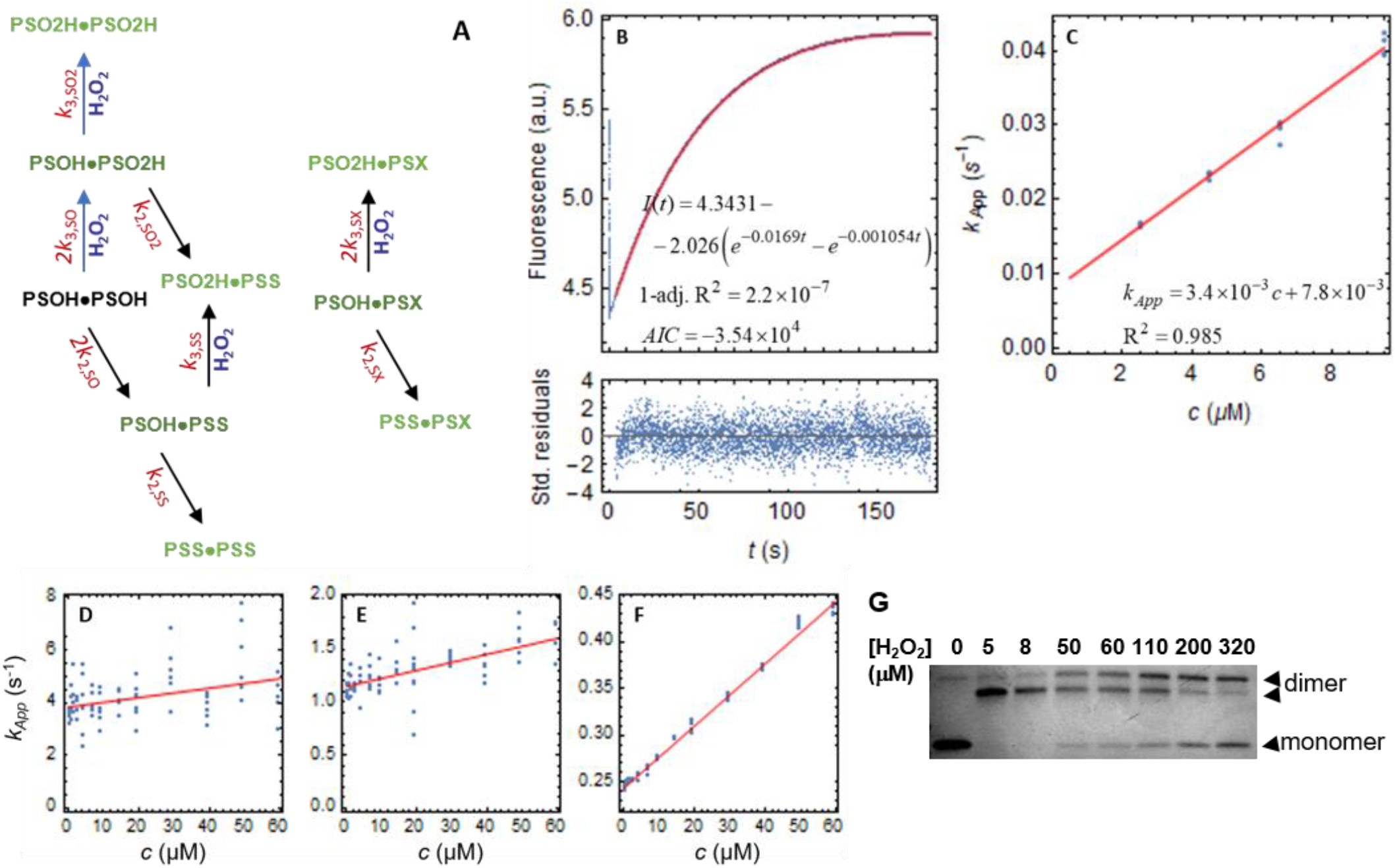
Analysis of fluorescence recovery curves for Prdx2 treated with excess H_2_O_2_ for evidence of cooperativity in hyperoxidation. (**A**) Reaction scheme for modelling the resolution of the sulfenic acids formed upon treatment of Prdx2 with supra-stoichiometric H_2_O_2_ concentrations. For the C_R_ mutant, the scheme simplifies as reactions involving the PSS forms cannot occur. Analysis of the resolution in WT Prdx2 required considering the fate of the sulfenic acids formed in anomalous dimers (right). Lighter shades of the symbols indicate higher fluorescence quantum yields, the precise equivalence of the quantum yields of the species in the same shade not being implied. (**B**) Example of full trace for 0.5 μM C172S Prdx2 treated with 3.0 μM H_2_O_2_ (blue) and bi-exponential fit to the ascending part (red). (**C**) H_2_O_2_-concentration dependence of the characteristic constant associated with a negative pre-exponential coefficient. *c* is the excess H_2_O_2_ over Prdx2 C_P_ sites. Blue, real data; red, best-fit curves. See SM2.2.4 for details of model and analysis. (**D, E, F**) H_2_O_2_ concentration dependence of the best-fit characteristic constants obtained for the Trp fluorescence recovery time course following treatment of 0.5 μM WT reduced Prdx2 with supra-stoichiometric H_2_O_2_ concentrations. *c* is the excess H_2_O_2_ over Prdx2 C_P_ sites. Red, best-fit lines. See Table 1 for best-fit values. Despite the large dispersion of the points in D the slope is statistically significant (p<0.002). (**G**) Hyperoxidation of Prdx2 (5 μM) analyzed by non-reducing SDS PAGE as in Fig 2. With stoichiometric H_2_O_2_, all the thiol groups are consumed and Prdx2 runs as lower two disulfide band. At higher concentrations the upper dimer band corresponds to one disulfide and one sulfinic acid, and the monomer band to the sulfinic acid.

We next measured the overall rate constant for sulfenic acid oxidation for WT Prdx2. Addition of H_2_O_2_ to reduced WT Prdx2 causes a rapid drop in fluorescence, followed by slower recovery. At low H_2_O_2_ concentrations, recovery is independent of H_2_O_2_ and relates to condensation to the disulfide (8,9). At higher concentrations, an H_2_O_2_-dependent component attributable to hyperoxidation is apparent (Fig S6 and (14)). This linear relationship gives an intercept of 0.3 s^-1^ (consistent with values previously determined for condensation using stopped flow (9,15)) and the slope corresponds to a rate constant for hyperoxidation of 4160 M^-1^s^-1^. We previously determined the hyperoxidation rate constants using catalase competition and MS analysis and obtained a similar values of approximately 6000 M^-1^s^-1^ (5).^1^

Further analysis of the fluorescence data using the approach in SM2.2.1 revealed that at all H_2_O_2_ concentrations tested, the fluorescence recovery was tri-exponential (Fig S8). The three characteristic constants increased linearly with the H_2_O_2_ concentration, with the slopes and intercepts at the origin shown in Table 1, experiment c (Fig 7D-F). We interpreted these results according to the scheme in Fig 7A with the intercepts corresponding to *k*_2_ for the different species and the slopes to *k*_3_. This creates a complex system with up to four exponential components, attributable to condensation and sulfinylation of the different sulfenic acid intermediates (SM2.2.4). The analysis of the data in light of this model and of the results previously obtained for hyperoxidation of the C172S mutant and for condensation in WT Prdx2 oxidized by stoichiometric H_2_O_2_ is explained in SM2.2.4. It led to the attribution of the above-mentioned slopes and intercepts to *k*_3,SS_, 2 *k*_3,SO_, *k*_3,SO2_, *k*_3,SX_ and *k*_2,SS_, 2 *k*_2,SO_, *k*_2,SO2_, *k*_2,SX_ (respectively) as shown in Table 2. The *k*_2_ values are remarkably similar to those obtained in the previous section. In turn, the values of *k*_3,SS_, *k*_3,SO_ and *k*_3,SO2_ are not significantly different from each other nor from the value obtained for the C_R_ mutant, which reinforces the conclusion that sulfinylation is non-cooperative. The higher value of *k*_3,SX_ suggests that sulfinylation of the anomalous dimers is substantially faster, though the large error in this estimate should be noted.

**Table 2:** Parameters determined in this work. The identities *k*_3,SO2_=*k*_3,SO_ and *k*_2,SO2_=*k*_2,SO_ were inferred from the fact that the fluorescence recovery associated to the sulfinylation of C172S Prdx2 is mono-exponential and that for WT Prdx2 is tri- and not tetra-exponential (see SM2.2.4 for details). *s*_y_=*k*_3_/*k*_2_.

| $k_{i,y}$ | | SH | SO | SS | SO2 | SX |
| --- | --- | --- | --- | --- | --- | --- |
| **1** | | $k_{1,\text{SH}}$ | (2.2±0.1)$\times k_{1,\text{SH}}$ | (1.25±0.07)$\times k_{1,\text{SH}}$ | - | - |
| **2** ($\text{s}^{-1}$) | a | - | 0.94±0.01 | 0.396±0.004 | - | 7.9±0.7 |
| | b | - | 0.54±0.01 | 0.2445±0.0004 | - | 3.28±0.04 |
| | c | - | 0.58±0.01 | 0.2435±0.0009 | 0.58±0.01 | 3.8±0.1 |
| **3** ($\text{M}^{-1}\text{s}^{-1}$) | C172S | - | (3.4±0.1)$\times 10^3$ | - | (3.4±0.1)$\times 10^3$ | - |
| | WT | - | (3.8±0.6)$\times 10^3$ | (3.33±0.03)$\times 10^3$ | (3.8±0.6)$\times 10^3$ | (18.±6)$\times 10^3$ |
| $s_y$ ($\text{M}^{-1}$) | | - | (7.±1)$\times 10^3$ | (1.37±0.01)$\times 10^4$ | (7.±1)$\times 10^3$ | (5.±2)$\times 10^3$ |

Note that a, b and c refer to the experiments described in Table 1.
The values for $k_{2,\text{SO}}$ are half the values of the intermediate characteristic constant for Trp fluorescence recovery because each $\text{PSOH}\cdot\text{PSOH}$ dimer carries two sulfenic acids that can condense.

The lower condensation rate constant *k*_2,SS_ translates into approximately twice the hyperoxidation susceptibility (*s*_y_= *k*_3,y_/*k*_2,y_) of the PSOH·PSS dimers relative to PSOH·PSOH and PSOH·PSO_2_H ones (Table 2). This result was independently assessed through gel experiments. Addition of stoichiometric H_2_O_2_ to Prdx2 oxidizes all the reduced C_P_ residues to form the 2-disulfide (lower) dimer. Higher concentrations cause progressive formation of the upper dimer, which contains one disulfide and one sulfinic acid then the hyperoxidized monomer (3,5) (Fig 7G). The kinetic model used above provided analytical solutions for the relative fractions of monomer, 1DS dimer and 2DS dimer formed, which when fitted to the densitometric determinations at various H_2_O_2_ concentrations gave estimates consistent with *s*_SO2_ ≈ *s*_SO_ ≈ *s*_SS_/2 (SM2.2.4)

Altogether, the results in this section (summarized in Table 2) show that: (i) the sulfinylation rate constant (*k*_3_) at one site is approximately the same irrespective of the other site being in sulfenic, sulfinic or disulfide form, (ii) the condensation rate constant at one site is approximately the same irrespective of the other site being in sulfenic or sulfinic form.

### Modest positive cooperativity for disulfide reduction by dithiothreitol

We also addressed the question of whether Prdx2 reduction is cooperative. Although not a physiological reductant, dithiothreitol (DTT) is widely used to reduce peroxiredoxins and other thiol proteins. Therefore, we followed the kinetics of Prdx2 disulfide reduction by DTT by SDS-PAGE (Fig 8). Consistent with a stepwise reduction of the two disulfides, the density of the 1DS band increased for the first 40 min and then gradually decreased. These results were analysed by fitting the model generated from the two site reaction scheme in Fig S2A to the densitometric readings over time. Best-fit values are *k*_4,SS_ = 3.75+0.05 and *k*_4,SH_ = 6.48+0.09 M^-1^s^-1^ (analysis in SI2.1.3), indicating positive cooperativity as the latter rate constant is 1.73±0.03 fold higher than the former.

**Figure 8.**
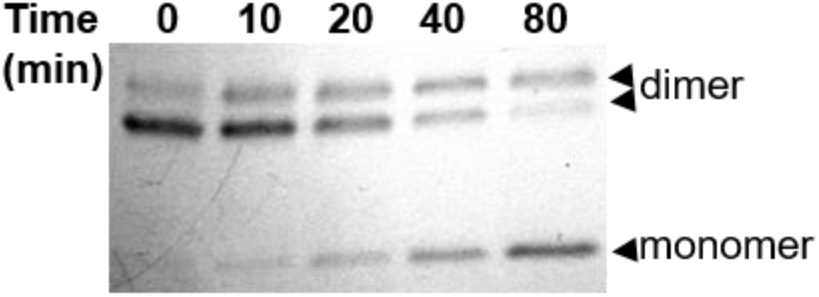
Reduction of Prdx2 disulfides by DTT. Oxidized Prdx2 (5 μM) was treated with 50 μM DTT. Samples were taken at the times shown, added to NEM and subjected to non-reducing SDS PAGE.

## Discussion

Our data indicate that there is cooperativity between the two active sites of the functional Prdx2 dimer in their reaction with H_2_O_2_, and kinetic modelling has enabled almost complete parameterization of the dimer-based reaction network. Separation of the reduced and disulfide forms by SDS PAGE has revealed that conversion of the reduced Cp thiol at the first site to a sulfenic acid increases the rate of oxidation of the second 2.3-fold. In contrast if the first site had time to condense to the disulfide before the second site was oxidized then both sites reacted similarly. In turn, negative cooperativity for the condensation of the sulfenic acid to disulfide was also identified through the analysis of Trp fluorescence changes using stopped flow. In this case, conversion of one sulfenic acid in PSOH.PSOH to a disulfide slowed the rate of condensation of the second to ~40%. The two rate constants obtained for condensation fall within the range of 0.24-1 s^-1^ determined by considering the two sites as indistinguishable (5,9,14). Best fit was obtained when a minor contribution of a third (faster) component of condensation reaction was included. We propose that this involves a species (X) in which one active site is in an apparently inactive form and which was present in small amounts in the Prdx preparations. Although the Trp fluorescence data were not sensitive enough to detect cooperativity in the first oxidation step, the two methodological approaches were consistent in showing no additional cooperativity at the hyperoxidation step. Similar rate constants of 3.4-3.8 × 10^3^ M^-1^s^-1^ were obtained regardless of whether the first site was PSOH, PSS or PSO_2_H. These compare with values of 2×10^3^ and 6×10^3^ M^-1^s^-1^ measured using stopped flow (14) and catalase competition (5,16) respectively.

An important consideration for interpreting these data is diffusion and whether apparent cooperativity could be due to H_2_O_2_ reacting with the closest Prdx2 molecule before mixing of the reactants was complete. This is not relevant for condensation, but is a potential issue for the very fast initial reaction with H_2_O_2_. We aimed to minimize this by using low micromolar concentrations, which taking a rate constant of ~10^8^ M^-1^s^-1^, can be calculated to give a reaction half life in the 5-10 ms range. Mixing therefore needs to be faster than this. Stopped flow mixing has a dead time of ~2 ms and enables fluorescence changes to be monitored within this time scale ((7–9) and Fig 5A). Mixing should therefore precede most of the reaction and not account for apparent cooperativity under these conditions. Furthermore, the similar cooperativity ratio obtained when the H_2_O_2_ was generated uniformly throughout the solution using pulse radiolysis supports the conclusion that it is a real phenomenon. However, the method of rapid mixing by vortexing that we employed appears to be insufficiently fast and we conclude that the higher apparent cooperativity ratio obtained with this method is due to oxidation occurring before mixing was complete. Nevertheless, this is a relevant consideration for experimental systems, which would typically give oxidation patterns as in Fig 2C. It is also physiologically relevant where limited diffusion of localized H_2_O_2_ in a cell is likely to cause oxidation of both active sites of peroxiredoxins in the vicinity rather than uniform oxidation of one site throughout the cell. A Prx2-dimer-based version of the reaction-diffusion model of cellular H_2_O_2_ signalling from (17), parameterized with the rate constants determined in this work, does indeed reveal enhanced positive cooperativity near the sites of H_2_O_2_ supply (results not shown).

As far as we are aware this is the first reported example of an enzyme combining positive and negative cooperativity in its catalytic cycle. An implication is that Prdx2 is in its most reactive form when one of its catalytic sites has been converted to the sulfenic acid. For this to happen, the second reaction must occur before the first sulfenic acid condenses, and a minimum H_2_O_2_ concentration is therefore required. The threshold above which this becomes more likely is given by [H_2_O_2_]=*k*_2,SH_/(*R*_1,SO_*k*_1,SH_). Assuming *k*_2,SH_ is in the range of *k*_2,SOH_ and *k*_2,SS_ and taking the k_2,SS_ value of 0.245/s, *R*_1,SO_=2 and *k*_1,SH_ = 100 μM^-1^s^-1^ gives a threshold of ~1.2 nM H_2_O_2_. This should be achievable with bolus addition or a high generation rate of H_2_O_2_, and within a cell may be favored where generation is localized and diffusion is restricted.

one expected consequence of the positive cooperativity for sulfenic acid formation and negative cooperativity for condensation would be to increase the lifetime of the PSS·PSOH form. The analytical approach taken by Selvaggio et al (18) indicates that any impact of cooperativity will vary depending on a number of factors, including Prdx and H_2_O_2_ concentrations and the activity of reductive mechanisms. Cooperativity at the first step is less likely to have an impact under conditions that maintain the Prxd in mainly a reduced form, and thus should be of little consequence when there is low-level oxidant production and adequate reductive capacity. It would have more impact where the steady state H_2_O_2_ concentration increases and more Prdx is oxidized. Considering the combination of positive cooperativity at the first step with the faster condensation of the double-sulfenic dimers, this could have a rate enhancing effect on the rate of H_2_O_2_ removal. The maximum possible enhancement is ≈2.2-fold and is most likely under situations when the main catalytic cycle is PSH·PSOH → PSOH·PSOH → PSOH·PSS → PSOH·PSH. This is the case in most human cell lines that have a high capacity for Prdx disulfide reduction (18). However, in cells such as erythrocytes with a low capacity for Prdx disulfide reduction, the sulfenic acid in PSOH·PSS would have time to condense before the disulfide is reduced, and negative cooperativity in condensation would decrease the rate of H_2_O_2_ removal under oxidative stress.

The influence of cooperativity on the preponderance of different redox intermediates could play a role in the regulation of redox signalling. There is mounting evidence that Prdxs can act as sensors that capture H_2_O_2_ then relay the oxidizing equivalents to other molecules (1,19–22). The most favored mechanism for this involves the sulfenic acid reacting with a thiol group on a partner protein. Specificity is likely to be influenced by the conformation and binding characteristics of the oxidized Prdx (23,24), and it is possible that different conformations of the sulfenic acid intermediates could influence the relay. Evidence for this remains to be explored. At H_2_O_2_ supply rates below cell’s capacity for reduction of Prdx1/2 disulfides redox relays are predicted to be spatially localized (17). Preliminary calculations with a reaction-diffusion model expanding on (17) indicate that the cooperativity characterized in this work has little effect on the spatial distribution of H_2_O_2_. However, cooperativity plus limited diffusion increases the concentration of PSOH·PSOH, PSOH·PSS, PSS·PSS, PSO_2_H·PSOH or PSO_2_H·PSS near H_2_O_2_ supply sites and strengthen the localization of redox relays mediated by these species. This effect may be further enhanced by the acidic environment near the membrane, which decreases all the condensation rates (Fig 6A) and strengthens cooperativity (Fig 6B).

Our findings of cooperativity imply that modification of one active site of Prdx2 influences the conformation of the other. Furthermore, to affect the initial reaction of C_P_ with H_2_O_2_, this must occur within a millisecond time scale. Typical 2-Cys Prdxs in their reduced form exist as decamers (or dodecamers) consisting of 5 (or 6) functional homodimeric units which are all present in their fully folded (FF) form (25–28). On complete oxidation to the disulfide the crystal structure remains decameric but the protein adopts a locally unfolded (LU) conformation in which C_R_ moves within disulfide bonding distance of C_P_ (25). Little attention has been given to asymmetry between sites, although there is suggestive evidence from molecular dynamic calculations (29), and crystal structures where the electron density is clear in one site but disordered in the other have been described (26,27). We do not have information on what changes might give rise to cooperativity. However, it seems unlikely that it relates to the FF → LU transformation. If this were the case, conversion of one site to the disulfide, which adopts the LU conformation, would be expected to increase the reactivity of the other.

With respect to the cooperativity in condensation, the characterized pH dependence of the characteristic constants does not support a mechanism mediated by a modulation of the pK_a_ of the C_P_-SOH or the C_R_-SH groups, as these appear to be similar for PSOH·PSS, PSOH·PSOH and PSOH·PSX dimers (Table S2). Instead, the rate constants for condensation between C_P_-SOH and C_R_-S^-^, C_P_-SO^-1^ and C_R_-S^-^, and C_P_-SOH and C_R_-SH all are 2.3-to 5.6-fold higher in PSOH·PSOH than in PSOH·PSS dimers. This is consistent with a scenario where the redox state of the second site induces a conformational change that influences the rate of the FF → LU transition (30) or the FF ↔ LU (quasi-) equilibrium (15) preceding and partially limiting the condensation reactions proper.

In conclusion, our kinetic studies have revealed that the active sites within the Prdx2 dimer do not behave independently. The conformational changes responsible for one site influencing the other are yet to be established and the effects are reasonably modest. Nevertheless, these findings identify a previously unrecognized molecular feature of Prdx2 that has potential for physiological consequences that warrant further investigation

## Acknowledgements

We are grateful to Dr Paul Pace and Dr Litiele Cezar da Cruzfor recombinant protein preparations and additional technical support.

## FOOTNOTES

This work was financed by the New Zealand Marsden Fund; by Fundação de Amparoa Pesquisa do Estado de São Paulo (FAPSEP) projects CEPID-Redoxoma 2013/07937-8 and Young Investigator 2018/14898-2 (Brazil); by the European Regional Development Fund, through COMPETE2020-Operational Program for Competitiveness and Internationalization, and Portuguese funds via FCT-Fundação para a Ciência e a Tecnologia, under project UID/NEU/04539/2019. L.S. is a postdoc funded by FAPESP 2017/12312-8.

The abbreviations used are: CP, peroxidatic Cys; CR, resolving Cys; DS, disulphide; DTT, dithiothreitol; FF, fully folded; LU, locally unfolded; Prdx, peroxiredoxin.

1 As described in 16. Peskin, A. V., Pace, P. E., and Winterbourn, C. C. (2019) Enhanced hyperoxidation of peroxiredoxin 2 and peroxiredoxin 3 in the presence of bicarbonate/CO2. *Free Radic. Biol. Med*. 145, 1-7, these have been corrected to half the values originally reported, due to an error in the calibration against catalase reactivity.

