## Supporting Information for "Intra-dimer cooperativity between the active site cysteines during the oxidation of peroxiredoxin 2"

### 1 Symbols list

| Symbol | Meaning |
|---|---|
| $B$ | Instrumental baseline fluorescence reading |
| $c$ | Excess $H_2O_2$ concentration over available Prdx2 $C_P$ thiolates |
| $f_0$ | Initial fraction of Prdx2 dimers with no disulfide bonds |
| $f_1$ | Initial fraction of Prdx2 dimers with one disulfide bond |
| $f_2$ | Initial fraction of Prdx2 dimers with two disulfide bonds |
| $f_{0,\infty}$ | Final fraction of Prdx2 dimers with no disulfide bonds |
| $f_{1,\infty}$ | Final fraction of Prdx2 dimers with one disulfide bond |
| $f_{2,\infty}$ | Final fraction of Prdx2 dimers with two disulfide bonds |
| $f_X$ | Fraction of Prdx2 dimers that cannot be oxidized to the 2-disulfide form by stoichiometric $H_2O_2$ addition (anomalous dimers) |
| $f_{0X}$ | Fraction of 0-disulfide dimers that are anomalous |
| $f_{1X}$ | Fraction of 1-disulfide dimers that are anomalous |
| $\Phi_{x,y}$ | Fluorescence quantum yield of dimers with $C_P$ in oxidation states *x*,*y*, where *x* and *y* can be:<br>SH, $S^-$<br>SOH, $SOH/SO^-$<br>SO2, $SO_2H/SO_2^-$<br>SX, unknown inactive state |
| $g_{SO}$ | Ratio between the rate constant for condensation of each sulfenic acid in a PSOH·PSOH dimer and that in a PSOH·PSS dimer |
| $G$ | Instrumental gain |
| $I$ | Fluorescence intensity signal |
| $k_{1,y}$ | Rate constant for sulfenylation<br>*y*= redox state of the second peroxidatic Cys, with meaning as above. |
| $k_{2,y}$ | Rate constant for condensation |
| $k_{2A}$ | Rate constant for condensation between a $C_P$-SOH and a $C_R$-$S^-$ |
| $k_{2B}$ | Rate constant for condensation between a $C_P$-$SO^-$ and a $C_R$-$S^-$ |
| $k_{2C}$ | Rate constant for condensation between a $C_P$-SOH and a $C_R$-SH |
| $k_{3,y}$ | Rate constant for sulfinylation |
| $k_{4,y}$ | Rate constant for disulfide reduction by DTT |
| $k^*_{1,y}$ | Pseudo-first-order rate constant for sulfenylation |

| | |
|---|---|
| $k^*_{3,y}$ | Pseudo-first-order rate constant for sulfinylation |
| $k_{Bl}$ | Rate constant for bleaching of the sulfinic acids |
| $k_C$ | Rate constant for condensation |
| $k_{Ia}$, $k_{Ib}$ | Rate constants for isomerisation between distinct Prdx2 conformations |
| $k_{Sa}$, $k_{Sb}$ | Rate constants for sulfinylation of distinct Prdx2 conformations |
| $p_0$ | Fraction of Prdx2 dimers in fully reduced form |
| $p_1$ | Fraction of Prdx2 dimers in mono-sulfenic forms |
| *Prdx2* | Concentration of Prdx2 dimers |
| $\theta_y$ | Scaled quantum yield increase associated to the condensation of a sulfenic acid in a dimer whose second $C_P$ is in oxidation state $y$ |
| $r_y$ | Resistance to sulfinylation of a sulfenic acid in a dimer whose second $C_P$ is in oxidation state $y$ |
| $R_{1,y}$ | Ratio $k_{1,y}/ k_{1,SH}$ |
| $R_{4,y}$ | Ratio $k_{4,SH}/ k_{4,SS}$ |
| $s$ | $H_2O_2$:Prdx2-monomer stoichiometry |
| $s_y$ | Susceptibility to sulfinylation of a sulfenic acid in a dimer whose second $C_P$ is in oxidation state $y$ |
| $t$ | Time |
| $v$ | Rate of $H_2O_2$ generation |
| $w_y$ | Scaled pre-exponential coefficient (pre-exponential weight) associated with $k_{2,y}$ or $k_{3,y}$ |

### 2 Data analysis

#### 2.1 Gel-based experiments

In all experiments, the proportions of monomer, dimer with one disulfide and dimer with two disulfides were measured by densitometry of Coomassie-stained non-reducing gels. Analysis of these data drew on the "overarching" dimer-based model for Prdx2 kinetics that was presented in ref. (1), in which the reactions of each active site are considered separately (Figure 1, main text). Consideration of known parameters and of the design of each experiment then prompted pertinent simplifications, which are explained and justified below and were used to fit the data.

Unless otherwise noted, all the numeric procedures were carried out in *Mathematica*™ v. 12.0.0.0, using the following functions and settings. Fits were done using the NonLinearModelFit function with the setting Method → "PrincipalAxis", or MaxIterations → 1000 when fits with the previous setting did not succeed. Systems of ordinary differential equations embodying the models were numerically integrated using the function ParametricNDSolveValue with the setting PrecisionGoal → 32.

##### 2.1.1 Slow $H_2O_2$ generation experiments to determine $R_{1,SS}$

In experiments where continuous slow generation of $H_2O_2$ ($v$=0.87 – 3.8 nM s$^{-1}$) was used with Prdx2 concentrations ≈10 µM, a quasi-steady-state $H_2O_2$ concentration $H_2O_{2,0} = \frac{v}{2k_{1,SH}Prdx2}$ initially establishes. The pseudo-first-order rate constant for sulfenylation of the initially formed PSOH·PSH dimers is thus $k^*_{1,SO} = R_{1,SO}k_{1,SH}H_2O_{2,0} = \frac{R_{1,SO}}{2}\frac{v}{Prdx2} < 5\frac{0.0038\,\mu M\,s^{-1}}{10\,\mu M} < 0.002\,s^{-1}$, assuming that $R_{1,SO} < 10$. (Here, $R_{1,SO} = k_{1,SO}/k_{1,SH}$ expresses how much faster the sulfenylation of the peroxidatic thiolate in a PSH·PSOH dimer is, relative to each of those in a PSH·PSH dimer.) Likewise, $k^*_{3,SO} = k_{3,SO}\frac{v}{k_{1,SO}Prdx2} < 10^{-2}\,\mu M^{-1}s^{-1}\frac{0.005\,\mu M\,s^{-1}}{10\,\mu M^{-1}s^{-1}10\,\mu M} = 5\times10^{-7}\,s^{-1} \ll k_{2,SO}$. Therefore, under these conditions virtually all the PSOH·PSH dimers form PSS·PSH. Both the contribution of the pathway PSH·PSH → PSOH·PSH → PSOH·PSOH → PSS·PSOH → PSS·PSS and sulfinylation reactions can thus be neglected (Figure 4A). Moreover, formation of PSS·PSH and PSS·PSS from PSH·PSH and PSS·PSH (respectively) is strongly ratelimited by the sulfenylation steps, rather than by the condensation steps. Accordingly, <0.25% of Prdx2 is in sulfenic form at any time: considering that all the $v < 5\,\text{nM s}^{-1}$ $H_2O_2$ generation forms sulfenic acids that condense at a ≥0.2 $s^{-1}$ rate constant, the concentration of sulfenic acids is $< \frac{5\,\text{nM s}^{-1}}{0.2\ \text{s}^{-1}} = 25\,\text{nM}$. Therefore, their possible reactivity with the NEM (2) used to stop the reaction will not affect the results.

$H_2O_2$ builds up as oxidation progresses and the pseudo-first-order rate constants above increase proportionally. However, the >100-fold excess of $k_{2,\text{SO}}$ over the initial $k^*_{1,\text{SO}}$ warrants that condensation of the PSOH·PSH remains faster than the sulfenylation of the second site until >99% of the Prdx2 is oxidized.

Altogether, the considerations above support using the following simplified model (**Model 1**) to fit the results from these experiments:

$$\begin{aligned}
\frac{d\,H_2O_2}{dt} &= v - 2k_{1,\text{SH}} PSH{\bullet}PSH\; H_2O_2 - R_{1,\text{SS}} k_{1,\text{SH}} PSS{\bullet}PSH\; H_2O_2 \\
\frac{d\,PSH{\bullet}PSH}{dt} &= -2k_{1,\text{SH}} PSH{\bullet}PSH\; H_2O_2 \\
\frac{d\,PSS{\bullet}PSH}{dt} &= 2k_{1,\text{SH}} PSH{\bullet}PSH\; H_2O_2 - R_{1,\text{SS}} k_{1,\text{SH}} PSS{\bullet}PSH\; H_2O_2 \\
\frac{d\,PSS{\bullet}PSS}{dt} &= R_{1,\text{SS}} k_{1,\text{SH}} PSS{\bullet}PSH\; H_2O_2
\end{aligned} \tag{1}$$

Here, $R_{1,\text{SS}} = k_{1,\text{SS}} / k_{1,\text{SH}}$ expresses how much faster the sulfenylation of the peroxidatic thiolate in a PSH·PSS dimer is, relative to each of those in a PSH·PSH dimer. The other symbols' meanings are as in Figure 1. At low $H_2O_2$ generation rates virtually all the Prdx2 was eventually oxidized to the 2-disulfide form. Accordingly, 1-disulfide dimers initially present in the Prdx2 samples (fraction $f_1$ of all the dimers) were treated as normal PSS·PSH, and the initial conditions were thus:

$$H_2O_2(0) = 0,\ PSH{\bullet}PSH(0) = (1 - f_1)Prdx2,\ PSS{\bullet}PSH(0) = f_1 Prdx2,\ PSS{\bullet}PSS(0) = 0. \tag{2}$$

In order to estimate $R_{1,\text{SS}}$, the dimer fractions obtained by numerical integration of equations (1) at the various observation times were fitted to the densitometrically determined ones as in the experiment above. Both $R_{1,\text{SS}}$ and $v$ were left as adjustable parameters so as to minimize the impact of inaccuracies in the determination of $v$. Because $k_{1,\mathrm{SH}}$ multiplies all terms in equations (1), uncertainties in the value of this parameter are fully absorbed in the best-fit estimate of $v$ and have virtually no effect on the best-fit values of $R_{1,\mathrm{SS}}$.

##### 2.1.2 End-point experiments with sub-stoichiometric bolus $H_2O_2$ additions to determine $R_{1,SO}$.

The model to fit these experiments has to deal with the following two facts about the reduced Prdx2 preparations used in the experiments. First, some of them contained a non-negligible initial fraction of 1-disulfide dimers ( $f_1 \approx 20\%$). Second, in some cases a fraction ($f_X$) of the dimers could not be oxidized to the 2-disulfide form by stoichiometric $H_2O_2$ addition.[1]

At the µM-range (sub-stoichiometric) $H_2O_2$ concentrations used, $\frac{k_{2,y}}{k_{1,y} H_2O_2} < 0.1$, $\frac{k_{3,y} H_2O_2}{k_{2,y}} < 0.01$, ($y = \mathrm{SO,SS,SO2}$) hold. (Here, $y$ represents the oxidation state of the second active site, in accordance with the convention in Figure 1 of the main text.) Therefore, all the active site thiols will be oxidized before any disulfides are formed and both the contribution of the pathway PSH·PSH → PSH·PSOH → PSH·PSS → PSOH·PSS → PSS·PSS for the formation of the 2-disulfide dimers, and sulfinylation reactions, can be neglected. Moreover, after all the $H_2O_2$ is consumed, all the PSOH·PSOH is committed to form PSS·PSS, and all the PSH·PSOH is committed to form PSH·PSS. One must also consider the possibility that the presence of the anomalous sites (denoted as PSX in Figure S 1 and in subsequent equations) influences the properties of the second site. The reaction scheme thus simplifies to that in Figure S 1 and the system of equations used as basis to fit the results of these experiments was, accordingly:

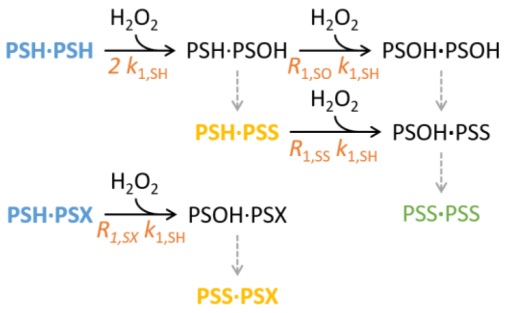

*Figure S 1. Simplified reaction scheme considered for fitting the end-point concentrations of monomer, 1- and 2-disulfide dimers. The slow reactions indicated by dashed arrows were not explicitly considered in the mathematical model. All the sulfenic acids are assumed to undergo condensation once all the $H_2O_2$ is consumed. The initially present species are represented in bold type. The stable species composing the monomer, 1-disulfide dimer and 2-disulfide dimer bands in non-reducing gels are represented in blue, yellow and green, respectively. The rate constants for the sulfenylation steps are represented in salmon. This reaction scheme is similar to that in Figure 4B except for the addition of the reactions involving the anomalous dimers.*

[1] Henceforth, these apparently not fully oxidizable dimers will be denoted as “anomalous”.

$$\frac{d\,H_2O_2}{dt} = -2k_{1,\mathrm{SH}} PSH{\bullet}PSH\; H_2O_2 - R_{1,\mathrm{SO}} k_{1,\mathrm{SH}} PSH{\bullet}PSOH\; H_2O_2$$
$$- R_{1,\mathrm{SS}} k_{1,\mathrm{SH}} PSH{\bullet}PSS\; H_2O_2 - R_{1,\mathrm{SS}} k_{1,\mathrm{SH}} PSH{\bullet}PSX\; H_2O_2$$
$$\frac{d\,PSH{\bullet}PSH}{dt} = -2k_{1,\mathrm{SH}} PSH{\bullet}PSH\; H_2O_2$$
$$\frac{d\,PSH{\bullet}PSOH}{dt} = 2k_{1,\mathrm{SH}} PSH{\bullet}PSH\; H_2O_2 - R_{1,\mathrm{SO}} k_{1,\mathrm{SH}} PSH{\bullet}PSOH\; H_2O_2$$
$$\frac{d\,PSOH{\bullet}PSOH}{dt} = R_{1,\mathrm{SO}} k_{1,\mathrm{SH}} PSH{\bullet}PSOH\; H_2O_2$$
$$\frac{d\,PSH{\bullet}PSS}{dt} = -R_{1,\mathrm{SS}} k_{1,\mathrm{SH}} PSH{\bullet}PSS\; H_2O_2$$
$$\frac{d\,PSOH{\bullet}PSS}{dt} = R_{1,\mathrm{SS}} k_{1,\mathrm{SH}} PSH{\bullet}PSS\; H_2O_2$$
$$\frac{d\,PSH{\bullet}PSX}{dt} = -R_{1,\mathrm{SX}} k_{1,\mathrm{SH}} PSH{\bullet}PSX\; H_2O_2$$
$$\frac{d\,PSOH{\bullet}PSX}{dt} = R_{1,\mathrm{SX}} k_{1,\mathrm{SH}} PSH{\bullet}PSX\; H_2O_2 \qquad (3)$$

with initial conditions:

$$H_2O_2(0) = H_2O_{2,0}$$
$$PSH{\bullet}PSH(0) = (1 - f_X - (1 - f_{1\mathrm{X}})\, f_1) Prdx2$$
$$PSH{\bullet}PSOH(0) = 0$$
$$PSOH{\bullet}PSOH(0) = 0$$
$$PSH{\bullet}PSS(0) = (1 - f_{1\mathrm{X}})\, f_1 Prdx2 \qquad (4)$$
$$PSOH{\bullet}PSS(0) = 0$$
$$PSH{\bullet}PSX(0) = f_X - f_{1\mathrm{X}}\, f_1 Prdx2$$
$$PSOH{\bullet}PSX(0) = f_{1\mathrm{X}}\, f_1 Prdx2$$

Here, $f_{1\mathrm{X}}$ and *Prdx2* stand for the fraction of the initial 1-disulfide dimers that are anomalous and for the total concentration of Prdx2 *dimeric units* (*i.e.*, half the total concentration of monomers), respectively. In the last lines of expressions (3) and (4) the PSOH·PSX was considered equivalent to PSS·PSX for simplicity.

The relative densities of the monomer, 1-disulfide dimer and 2-disulfide dimer gel bands correspond respectively to the fractions

$$f_{0,\infty}=\frac{PSH\bullet PSH+PSH\bullet PSX}{Prdx2},$$
$$f_{1,\infty}=\frac{PSH\bullet PSOH+PSH\bullet PSS+PSS\bullet PSX}{Prdx2}, \qquad (5)$$
$$f_{2,\infty}=\frac{PSOH\bullet PSOH}{Prdx2},$$

(respectively) obtained by numerically integrating equations (3) from $t$=0 to $t$= 60 s, which gives sufficient time to consume all the $H_2O_2$. Formally, the functions obtained by replacing into equations (5) the $t\to\infty$ limit of the solutions of the system of equations (3) with the initial conditions (4) constitute **Model 2**. Because in this approximate model $k_{1,\mathrm{SH}}$ appears as a common factor in the right hand side of all equations in (3), it affects how fast $f_{0,\infty}$, $f_{1,\infty}$ and $f_{2,\infty}$ are approached but not the fractions themselves. Therefore, the value chosen for this parameter does not influence the best-fit estimates for $R_{1,\mathrm{SO}}$. It was assumed that $R_{1,\mathrm{SS}}=1$, as determined from the experiments with slow $H_2O_2$ generation (Section 2.1.1).

Model 2 has up to six adjustable parameters — $R_{1,\mathrm{SO}}$, $R_{1,\mathrm{SX}}$, $f_1$, $f_X$, $f_{1\mathrm{X}}$ and $Prdx2$ —, only some of which identifiable in practice (*i.e.*, tightly constrainable by the fits). Moreover, in some experiments $f_1$, $f_X$ and/or $f_{1\mathrm{X}}$ are negligible or have little influence on the best-fit estimates of $R_{1,\mathrm{SO}}$. To cope with these facts and ensure that the most parsimonious possible statistical model is used to fit each data set we adopted the following strategy. We used Model 2 as a template to generate a family of simpler statistical models by alternatively:

1. setting $R_{1,\mathrm{SX}}$ to 1 or $R_{1,\mathrm{SO}}$, corresponding to assuming that the second $C_P$-$S^-$ in an anomalous dimer is as reactive as that of a PSH·PSS or a PSH·PSH, respectively;
2. setting $f_1$ to the relative density of the 1-disulfide gel band obtained with no $H_2O_2$ addition or letting it adjust to account for the experimental uncertainty of this determination;
3. setting $f_X$ to 0 or letting it adjust;
4. setting $f_{1\mathrm{X}}$ to 0, $f_X$ or $f_{1,\max}\doteq\min\left(f_X/\max(0.01,f_1),1\right)$, respectively corresponding to assuming that before $H_2O_2$ addition the anomalous dimers were all in the monomer

fraction, distributed with equal probability between the monomer and 1-disulfide fractions, or distributed with the strongest preference to the 1-disulfide dimers;

5. setting *Prdx2* to the nominal concentration used in each experiment, or leaving it adjustable to minimize the impact of inaccuracies in the determination of low-µM Prdx2 and $H_2O_2$ concentrations.

*Table S 1. Set of statistical models used to fit band density data sets. See text for details. a, adjustable; s, fixed.*

| ***Prdx2*** | $\boldsymbol{f_1}$ | $\boldsymbol{f_X}$ | $\boldsymbol{f_{1X}}$ | $\boldsymbol{R_{1,SX}}$ |
|---|---|---|---|---|
| a | a | 0 | 0 | - |
| a | a | a | 0 | 1 |
| a | a | a | 0 | $R_{1,SO}$ |
| a | a | a | $f_X$ | 1 |
| a | a | a | $f_X$ | $R_{1,SO}$ |
| a | a | a | $f_{1,max}$ | - |
| a | s | 0 | 0 | - |
| a | s | a | 0 | 1 |
| a | s | a | 0 | $R_{1,SO}$ |
| a | s | a | $f_X$ | 1 |
| a | s | a | $f_X$ | $R_{1,SO}$ |
| a | s | a | $f_{1,max}$ | - |
| s | a | 0 | 0 | - |
| s | a | a | 0 | 1 |
| s | a | a | 0 | $R_{1,SO}$ |
| s | a | a | $f_X$ | 1 |
| s | a | a | $f_X$ | $R_{1,SO}$ |
| s | a | a | $f_{1,max}$ | - |
| s | s | 0 | 0 | - |
| s | s | a | 0 | 1 |
| s | s | a | 0 | $R_{1,SO}$ |
| s | s | a | $f_X$ | 1 |
| s | s | a | $f_X$ | $R_{1,SO}$ |
| s | s | a | $f_{1,max}$ | - |

Altogether, this generated the 24 statistical models schematized in Table S 1. In order to estimate $R_{1,SO}$, the values of $f_{0,\infty}$, $f_{1,\infty}$ and $f_{2,\infty}$ computed by numerical integration of each of the 24 models above for the various sub- to equi-stoichiometric $H_2O_2$ bolus additions were fitted to the relative densities of the monomer, 1-disulfide dimer and 2-disulfide dimer bands. The models were ranked in inverse order of the respective fits Akaike Information Criterion (3) with small-sample correction (AICc). For each data set we selected among the statistical models that yielded best-fit estimates significantly different from 0 for all the adjustable parameters the one with the lowest AICc value. The $R_{1,SO}$ estimate and standard error for each experiment were obtained from the respective selected model.

In a few experiments, treatment of Prdx2 with stoichiometric or slightly supra-stoichiometric $H_2O_2$ left a residual monomer band, and in others the initial Prxd2 sample contained some 2-disulfide dimer. We treated these as non-reactive components. Accordingly, we subtracted the densities of the corresponding bands from the densities of the monomer or 2-disulfide dimer bands (respectively) for all the $H_2O_2$ concentrations and recalculated the relative densities of all the bands with reference to the discounted total.

We also used a variant of Model 2 explicitly considering condensation of all the sulfenic acids with a 0.4 $s^{-1}$ rate constant (intermediate among previous experimental determinations) as basis for generating a similar set of statistical models. However, these models in general yielded worse fits, or fits that were not significantly better and produced best-fit $R_{1,SO}$ estimates similar to those based in Model 2.

##### 2.1.3 Analysis of cooperativity in Prdx2 reduction by dithiothreitol

The time course of the reduction of Prdx2 disulfide dimers by excess dithiothreitol (DTT) can be described by the reaction scheme in Figure S 2A. Assuming that the concentration of reduced DTT remains approximately constant, this translates into the following system of equations:

$$\begin{aligned}\frac{d\,PSS\bullet PSS}{d\,t} &= -2k_{4,SS} DTT\; PSS\bullet PSS \\ \frac{d\,PSH\bullet PSS}{d\,t} &= 2k_{4,SS} DTT\; PSS\bullet PSS - R_{4,SH} k_{4,SS} DTT\; PSH\bullet PSS \\ \frac{d\,PSH\bullet PSH}{d\,t} &= R_{4,SH} k_{4,SS} DTT\; PSH\bullet PSS.\end{aligned} \tag{6}$$

Here, $k_{4,SS}$ is the rate constant for reduction of a disulfide in a PSS·PSS dimer, and $R_{4,SH}$ is the ratio between the rate constant $(k_{4,SH})$ for reduction of the disulfide in a PSH·PSS dimer and $k_{4,SS}$. The initial conditions are:

$$\begin{aligned}PSS\bullet PSS(0) &= (1 - f_0 - f_1) Prdx2 \\ PSH\bullet PSS(0) &= f_1\, Prdx2 \\ PSH\bullet PSH(0) &= f_0\, Prdx2.\end{aligned} \tag{7}$$

In turn, with these initial conditions the solution of system (6), expressed in terms of the relative fractions of 0-, 1- and 2-disulfide dimers, is:

$$\frac{PSS\bullet PSS(t)}{Prdx2}=1+\frac{\frac{R_{4,\mathrm{SH}}}{2}}{1-\frac{R_{4,\mathrm{SH}}}{2}}(1-f_0-f_1)e^{-2k_{4,\mathrm{SS}}DTT\,t}-\frac{1-f_0-\frac{R_{4,\mathrm{SH}}}{2}f_1}{1-\frac{R_{4,\mathrm{SH}}}{2}}e^{-R_{4,\mathrm{SH}}k_{4,\mathrm{SS}}DTT\,t}$$

$$\frac{PSH\bullet PSS(t)}{Prdx2}=-\frac{1-f_0-f_1}{1-\frac{R_{4,\mathrm{SH}}}{2}}e^{-2k_{4,\mathrm{SS}}DTT\,t}+\frac{1-f_0-\frac{R_{4,\mathrm{SH}}}{2}f_1}{1-\frac{R_{4,\mathrm{SH}}}{2}}e^{-R_{4,\mathrm{SH}}k_{4,\mathrm{SS}}DTT\,t} \qquad (8)$$

$$\frac{PSH\bullet PSH(t)}{Prdx2}=(1-f_0-f_1)e^{-2k_{4,\mathrm{SS}}DTT\,t}$$

Equations (8) constitute **Model 3**.

This model yielded excellent fits to the time evolution of experimental relative dimer fractions for two independent experiments where 5 µM oxidized Prdx2 was reduced by 50 µM DTT at room temperature, with samples taken over 80 min (Figure S 2B,C). Both experiments yield similar best-fit estimates for $k_{4,\mathrm{SS}}$ and $R_{4,\mathrm{SH}}$, giving variance-weighted means $k_{4,\mathrm{SS}}$ = 3.75±0.05 $M^{-1}s^{-1}$ and $R_{4,\mathrm{SH}}$ = 1.73±0.03. The latter value reveals a modest positive cooperativity, such that the reduction of the first disulfide in a dimer facilitates the reduction of the second.

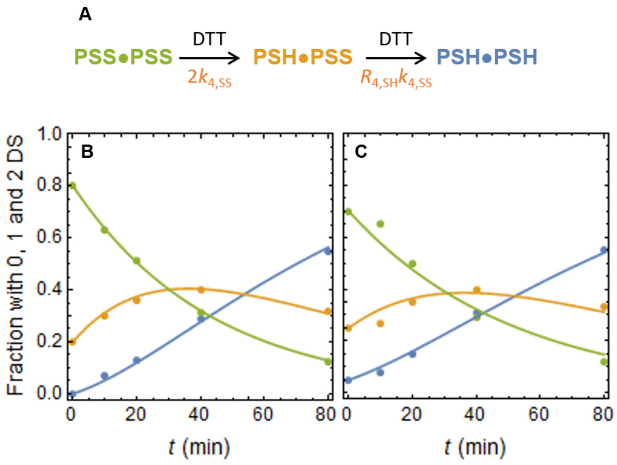

| | B | C |
|---|---|---|
| $R_{4,\mathrm{SH}}$ | 1.72±0.05 | 1.8±0.2 |
| $k_{4,\mathrm{SS}}$ | 3.82±0.07 $M^{-1}s^{-1}$ | 3.2±0.2 $M^{-1}s^{-1}$ |
| Adj. $R^2$ | 0.9992 | 0.994 |
| AICc | -89.0 | -56. |

*Figure S 2. Modeling of Prdx2 reduction by DTT. (A) Reaction scheme. Rate constants are indicated in salmon. (B,C) Fits of Model 3 (solid lines) to the evolution of the relative fractions of 0- (blue), 1- (yellow) and 2-disulfide (green) dimers, measured by densitometric analysis of SDS-PAGE gels. The table below the plots presents the best-fit parameters and goodness-of-fit statistics.*

#### 2.2 Analysis of Trp quenching experiments

Prdx2's Trp fluorescence intensity changes over several time scales upon $H_2O_2$ addition. It first decreases in a few ms to ~0.1 s, and then undergoes a multi-exponential recovery over ~10 s. The initial fluorescence quenching is associated to the binding of $H_2O_2$ and formation of the sulfenic acid, whereas the subsequent slow fluorescence recovery is associated to the condensation of the sulfenic acid and to its oxidation to a sulfinic acid (4–7). Because the two phases of the fluorescence change report on distinct molecular events we analysed them separately. The subsections below discuss these analyses.

##### 2.2.1 Procedure for analysis of the fluorescence recovery phase

###### *2.2.1.1 General workflow*

In order to capture both the fast and the slow changes following each stopped-flow injection, data were acquired at exponentially increasing times, with time steps and integration times ranging from 0.0125 ms to 46.4 ms. Handling the issues raised by the non-monotonicity, variable sampling time step and signal integration time, multi-exponentiality of the fluorescence recovery and bleaching, and then providing a mechanistic interpretation of the results required a relatively complex workflow (Figure S 3). The rationale for and details of each step in this workflow are explained in the next subsections.

##### 2.2.1.2 Data windowing

The fluorescence recovery time series were trimmed at both short and long times for different reasons. For a short time following the fluorescence minimum ($t = t_{min}$) the fluorescence intensity still reflects the relaxation of the quenching processes, as evident from the local upwards concavity. (An exponential fluorescence recovery traces a concave-down curve instead.) Inclusion of the period immediately following $t_{min}$ in the time series can substantially biase the estimation of the fastest characteristic constants of fluorescence recovery. In turn, visual inspection revealed that the fluorescence intensity time series were consistently concave-down beyond $t_{min} + 0.1$ s, and a systematic analysis revealed that optimal consistency in best-fit estimates among replicates and over added [$H_2O_2$] was achieved when the data points at $t < t_{min} + 0.2$ s were neglected. Therefore, we neglected the data points at $t < t_{min} + 0.2$ s, though considering $t = t_{min}$ as the starting time for fluorescence recovery. The observations that the residuals of mono-exponential fits are consistently negative at the $t_{min} + 0.2$ s starting points and that all the pre-exponential coefficients in the multi-exponential fits are negative indicate that this procedure effectively eliminates any significant influence of the dying-off of the fluorescence quenching phase.

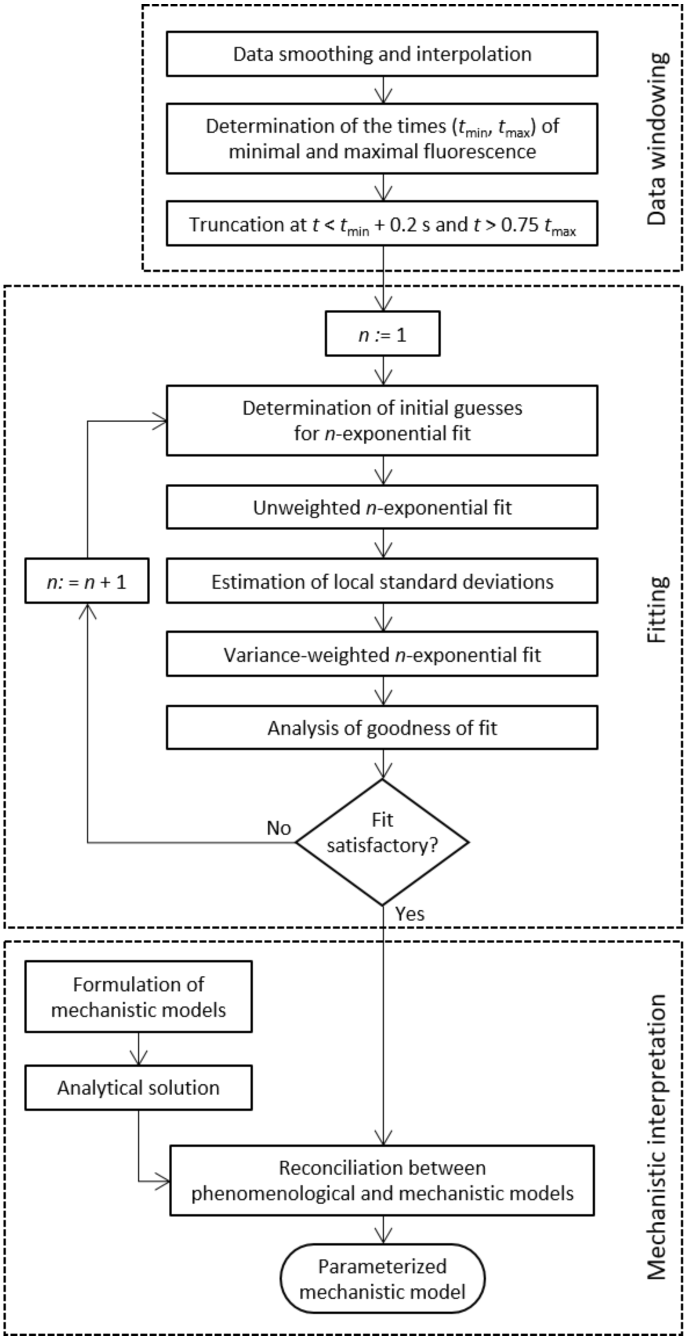

*Figure S 3. Trp fluorescence recovery data analysis workflow.*

In some replicates there was a barely noticeable decrease in fluorescence intensity at long times ($t > 15$ s), likely due to bleaching of the Trp residues (8). In order to minimize the influence of this decrease on the parameter estimates we determined the time of occurrence of the fluorescence maximum ($t_{max}$) and neglected the data for $t \geq 0.75\ t_{max}$. The observations that all the pre-exponential coefficients in the multi-exponential fits are negative and that the estimated parameters do not drastically change when the $t \geq 0.75\ t_{max}$ portion of the time series is included indicate that this procedure effectively eliminates any significant influence

The $t_{min}$ and $t_{max}$ values were determined from the time series as follows. We applied a low-pass filter with cut-off frequencies 0.03 (for $t_{min}$) or 0.05 (for $t_{max}$) to the time series (function LowpassFilter in *Mathematica*™ v. 12.0.0.0 (9)). The quality of the filtering was visually inspected. We then determined $t_{min}$ and $t_{max}$ by applying *Mathematica*™ functions NMinimize (with the constraint $0.002\ \text{s} < t \leq 0.14\ \text{s}$) and NMaximize (with $2\ \text{s} < t \leq 20\ \text{s}$), respectively, to third-order polynomial interpolations of the filtered data.

###### 2.2.1.3 Multi-exponential fitting

The multi-exponential fits to the windowed unfiltered time series were carried out through the following iterative procedure. Starting with mono-exponential [$n$= 1 in expression (9)] fits, fits of linear combinations of $n$+1 exponentials,

$$I(t) = b + \sum_{i=1}^{n+1} a_i e^{-k_i t} \tag{9}$$

were attempted when both the following criteria were satisfied: (i) The residuals of the fit to a linear combination of $n$ exponentials were not randomly distributed around 0 over time, and instead showed a clear “undulating” bias; (ii) the same pattern of residuals was consistently reproduced over replicates.

Linear combinations of $n$+1 exponentials were judged good statistical models of the data whenever the following four criteria were cumulatively satisfied:

1. The fits yielded plausible estimates and tight confidence intervals for all adjustable parameters.
2. The systematic bias in the fit residuals were attenuated or eliminated.

3. The Akaike Information Criterion for the $n$+1-exponentials fit is at least 12 units lower than that for the $n$-exponentials, the threshold to warrant that the former model has a higher likelihood at 99.75% ($p<0.0025$) significance.
4. The parameter estimates were consistent across replicates.

Importantly, this procedure determines the number of significant components in an objective way while avoiding overfitting.

An exponential time sampling scheme was used that implied longer signal integration times at late times than at early times. As a consequence, the time series for Trp fluorescence recovery had non-uniform error variances over the response variable (heteroscedasticity), which might bias parameter estimation. In order to correct for these biases, variance-weighted fits were carried out. Local standard deviations for this purpose were estimated through a linear regression of the absolute values of the residuals from the unweighted fits against the (unweighted) fitted values (10).

All the fits were carried out in *Mathematica*™ v. 12.0.0.0 using the function NonlinearModelFit with options set to ensure appropriate variance weighting (where pertinent) and full convergence of the error minimization (setting MaxIterations $\rightarrow$ 5000 unless otherwise stated).

Successful multi-exponential fitting of time series frequently requires good initial guesses of the best-fit parameters, which may be cumbersome to achieve by trial and error. Therefore, we implemented the CEF algorithm (11) in *Mathematica*™ and used it to obtain accurate initial guesses.

###### 2.2.1.4 Mechanistic interpretation

This part of the work addresses whether and to what extent the redox state of one active site in a Prdx2 dimer influences the rate constants for condensation and sulfinylation at the other site. The number and relative values of the characteristic constants and pre-exponential coefficients determined through the analysis described in the previous section, together with the knowledge of generic molecular properties of Prxd2, can guide mechanistic interpretations. However, complex data are often amenable to alternative interpretations. Our strategy to minimize this possibility draws on the following three principles:

First, we used a stepwise modular approach. Thus, we initially performed experiments with stoichiometric $H_2O_2$ addition to low Prdx2 concentrations to examine the kinetics of condensation under conditions where sulfinylation is negligible (Section 2.2.2). We then performed experiments with supra-stoichiometric $H_2O_2$ addition to a resolving Cys mutant Prdx2 (C172S Prdx2) to examine the kinetics of sulfinylation under conditions where condensation cannot occur (Section 2.2.3). Finally, we performed experiments with equi- and supra-stoichiometric $H_2O_2$ addition to Prdx2 to examine the kinetics of condensation and sulfinylation together (Section 2.2.4), drawing on the results from the previous experiments for proper interpretation.

Second, we used the most parsimonious mechanistic models possible to explain each data set, thus maximizing mathematical tractability and avoiding to the extent possible to deal with mechanistic uncertainties that the data itself does not allow to clarify.

Third, we designed experimental conditions such that all the relevant reactions have pseudo-first-order kinetics. This allowed the formulation of analytically solvable mechanistic models. Analytical solutions permitted relating the signs and relative values of the observational characteristic constants and pre-exponential coefficients to mechanistic features in a relatively straightforward manner. Additionally, the algebraic analysis of the solutions permitted finding combinations of the observational parameters that (i) are independent of the instrumental baseline and gains, and (ii) can be analytically expressed as function of combinations of mechanistic parameters that have both intuitive physical meaning and well-defined ranges of plausible values. These features proved crucial for the following verification.

Fourth, we reconciled mechanistic models to phenomenological models by verifying that the observational parameters are quantitatively consistent with all the theoretical characteristic constants and pre-exponential coefficients that were derived from the mechanistic models.

The alternative approach of directly fitting numerical mechanistic models to the data has several disadvantages. Namely, it yields less insight on the relationship between mechanistic parameters and features of the time series; it requires handling a larger number of adjustable parameters, which at best increases the uncertainty of the relevant estimates; it makes the fitting much more computationally demanding and less robust.

The approach described above led to a fully parameterized mechanistic (kinetic) model for the reaction of Prdx2 with equi- and supra-stoichiometric $H_2O_2$, considering all the intra-dimer allosteric effects between active sites that can occur under these conditions.

##### 2.2.2 Analysis of condensation of Prdx2 with stoichiometric $H_2O_2$ addition

These experiments were carried out under conditions where the pseudo-first-order rate constant for sulfenylation is >$10^3$-fold that for condensation, and the second-order rate constant is ~$10^4$-fold that for sulfinylation. Therefore, when $H_2O_2$ is added to Prdx2 in stoichiometric amounts both $C_P$ thiolates in a dimer are sulfenylated before significant condensation or sulfinylation can occur. It is thus a good approximation to consider that all the available Prdx2-$C_P$ thiolates are instantaneously sulfenylated. On the other hand, the Prdx2 samples contained a non-negligible initial fraction of 1-disulfide dimers $(f_1)$, and of anomalous dimers (fractions $f_{0X}$ and $f_{1X}$ of the 0- and 1-disulfide dimers, respectively).[2] Considering that the anomalous sites may influence the properties of the second site, we used the following equations to analyze the Trp fluorescence recovery upon treatment of Prdx2 with stoichiometric $H_2O_2$ (Figure S 4):

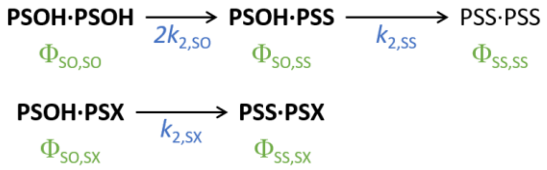

*Figure S 4. Reaction scheme considered for simulating the Trp fluorescence recovery experiments with stoichiometric bolus $H_2O_2$ addition to Prdx2. The species present immediately after sulfenylation are represented in bold type. The fluorescence quantum yields of each species are represented in green. The hydrogen peroxide concentration used is low enough to cause negligible hyperoxidation. The reaction scheme is identical to that in Figure 5E and repeated here to highlight the quantum yield symbols.*

$$
\begin{aligned}
\frac{d\,PSOH\bullet PSOH}{d\,t} &= -2k_{2,\mathrm{SO}}PSOH\bullet PSOH \\
\frac{d\,PSOHPSS}{d\,t} &= 2k_{2,\mathrm{SO}}PSOH\bullet PSOH - k_{2,\mathrm{SS}}PSOH\bullet PSS \\
\frac{d\,PSOH\bullet PSX}{d\,t} &= -k_{2,SX}\,PSOH\bullet PSX \\
\frac{d\,PSS\bullet PSS}{d\,t} &= k_{2,\mathrm{SS}}PSOH\bullet PSS \\
\frac{d\,PSS\bullet PSX}{d\,t} &= k_{2,\mathrm{SS}}PSOH\bullet PSX
\end{aligned}
\qquad (10)
$$

[2] We will henceforth denote by "0-disulfide" any dimers, anomalous or not, that dissociate to monomers in non-reducing gels.

with the initial conditions

$$\begin{aligned}
&PSOH\bullet PSOH(0) = (1-f_{0X})(1-f_1)Prdx2,\\
&PSOH\bullet PSS(0) = (1-f_{1X})\,f_1 Prdx2,\\
&PSOH\bullet PSX(0) = f_{0X}(1-f_1)Prdx2,\\
&PSS\bullet PSS(0) = 0,\\
&PSS\bullet PSX(0) = f_{1X}\,f_1 Prdx2.
\end{aligned} \tag{11}$$

Considering that the time evolution of the fluorescence intensity signal ($I$) is given by

$$\begin{aligned}
I(t) = B + G(&\Phi_{SO,SO} PSOH\bullet PSOH(t) + \Phi_{SO,SS} PSOH\bullet PSS(t) + \Phi_{SS,SS} PSS\bullet PSS(t) + \\
&+\Phi_{SO,SX} PSOH\bullet PSX(t) + \Phi_{SS,SX} PSS\bullet PSX(t)),
\end{aligned} \tag{12}$$

with $\Phi_{x,y}$ the fluorescence quantum yields of the various dimer forms, and replacing the solution of equations (10) one finds a solution in the form:

$$I(t) = b + c_{SX} e^{-k_{2,SX}t} + c_{SO} e^{-2k_{2,SO}t} + c_{SS} e^{-k_{2,SS}t} \tag{13}$$

with:

$$\begin{aligned}
&b = B + G\ Prdx2\left(\Phi_{SS,SS} + (f_{0X}(1-f_1) + f_{1X} f_1)(\Phi_{SS,SX} - \Phi_{SS,SS})\right),\\
&c_{SX} = -G\ Prdx2\ f_{0X}(1-f_1)(\Phi_{SS,SX} - \Phi_{SO,SX}),\\
&c_{SO} = -G\ Prdx2\ (1-f_{0X})(1-f_1)\left(\Phi_{SS,SS} - \Phi_{SO,SO} - \kappa(\Phi_{SO,SS} - \Phi_{SO,SO})\right),\\
&c_{SS} = -G\ Prdx2\left((1-f_{0X})(1-f_1)\kappa + (1-f_{1X})\,f_1\right)(\Phi_{SS,SS} - \Phi_{SO,SS}),\\
&\kappa = \frac{2k_{2,SO}}{2k_{2,SO} - k_{2,SS}}.
\end{aligned} \tag{14}$$

Equation (13) constitutes **Model 5**. Although the pre-exponential coefficients depend on the instrumental gain, absolute quantum yields and Prdx2 concentration, their relative contributions for the fluorescence change, $w_y = c_y / (c_{SX} + c_{SO} + c_{SS})$, do not and are thus easier to interpret with relation to the experimental results:

$$\begin{aligned}
w_{\mathrm{SX}} &= \frac{f_{0\mathrm{X}}(1-f_1)\theta_{\mathrm{SX}}}{D},\\
w_{\mathrm{SO}} &= \frac{(1-f_{0\mathrm{X}})(1-f_1)\left(1-\kappa\,\theta_{\mathrm{SO}}\right)}{D},\\
w_{\mathrm{SS}} &= \frac{\left((1-f_{0\mathrm{X}})(1-f_1)\kappa+(1-f_{1\mathrm{X}})f_1\right)(1-\theta_{\mathrm{SO}})}{D},\\
D &= 1-f_{1\mathrm{X}}f_1-f_{0\mathrm{X}}(1-f_1)(1-\theta_{\mathrm{SX}})-(1-f_{1\mathrm{X}})f_1\theta_{\mathrm{SO}},\\
\theta_{\mathrm{SX}} &= \frac{\Phi_{\mathrm{SS,SX}}-\Phi_{\mathrm{SO,SX}}}{\Phi_{\mathrm{SS,SS}}-\Phi_{\mathrm{SO,SO}}},\ \theta_{\mathrm{SO}} = \frac{\Phi_{\mathrm{SO,SS}}-\Phi_{\mathrm{SO,SO}}}{\Phi_{\mathrm{SS,SS}}-\Phi_{\mathrm{SO,SO}}}.
\end{aligned} \tag{15}$$

The analysis of equations (14) yields several useful insights on the interpretation of the experimental results. First, if none of the initially fully reduced dimers contains anomalous sites $(f_{0\mathrm{X}}=0)$ then $c_{\mathrm{SX}}=0,$ and thus the fluorescence recovery will be at most bi-exponential. Second, considering that in typical experiments $f_{0\mathrm{X}}<1$ and $f_1<1,$ $c_{\mathrm{SO}}$ will be null only if $k_{2,\mathrm{SS}}=k_{2,\mathrm{SO}}$ *and* $\Phi_{\mathrm{SS,SS}}-\Phi_{\mathrm{SO,SO}}=2(\Phi_{\mathrm{SS,SS}}-\Phi_{\mathrm{SO,SS}}).$ That is, only if condensation at one site influences *neither* the rate constant for nor the change in fluorescence quantum yield associated to condensation at the second site. (Bar the unlikely case of the effect on the quantum yield change exactly compensating the effect on the rate constant, $\frac{\Phi_{\mathrm{SS,SS}}-\Phi_{\mathrm{SO,SO}}}{\Phi_{\mathrm{SS,SS}}-\Phi_{\mathrm{SO,SS}}}=\kappa,\ \kappa\neq 2.$) Therefore, a tri-exponential Trp fluorescence recovery will reveal allosteric cross-talk between the two active sites. Moreover, if none of the characteristic constants is twice another one, then condensation at one site influences the rate constant for condensation at the other site.

We fitted multi-exponential models to Trp fluorescence recovery time courses from an experiment using approximately stoichiometric $H_2O_{2,0}$ according to the workflow described in Figure S 3. The initial fractions of 1-disulfide dimers $(f_1=0.075)$ and of anomalous dimers $(f_{0\mathrm{X}}(1-f_1)+f_{1\mathrm{X}}f_1=0.082)$ were determined by non-reducing SDS-PAGE of the Prdx2 in the outflow from the stopped-flow apparatus, the former with no $H_2O_2$ and the latter after addition of 1.6 µM $H_2O_2$ (seven replicate stopped-flow injections) to 1.3 µM reduced protein.

The Trp fluorescence recovery from the latter $H_2O_2$ addition was tri-exponential (example in Figure 5B-D), with all the pre-exponential coefficients negative, and the characteristic constants and pre-exponential weights in Table 1, experiment a. The fact that all the characteristic constants are different and none twice another one demonstrates that there is cooperativity in condensation, as per the theoretical analysis above. However, the determination of the nature and extent of that cooperativity hinges on the correct assignment of the characteristic constants above to $k_{2,\mathrm{SX}}$, $2k_{2,\mathrm{SO}}$ and $k_{2,\mathrm{SS}}$. Upon replacement into equations (15) and consideration of the experimentally determined values of $f_1$, of the total fraction of anomalous dimers $\left(f_X = f_{0\mathrm{X}}(1-f_1)+f_{1\mathrm{X}}f_1\right)$ and plausible ranges for $\theta_{\mathrm{SX}}$, $\theta_{\mathrm{SO}}$, $f_{0\mathrm{X}}$ and $f_{1\mathrm{X}}$ the correct assignments must yield ranges of $w_X$, $w_O$ and $w_C$ that are consistent with the experimental values of these weights. Our analysis of this consistency is based on the following considerations. First, the experimental values of $f_1$ and $f_X$ determine the relationship $f_{0\mathrm{X}} = 0.089 - 0.081 f_{1\mathrm{X}}$. Second, the fraction of anomalous 1-disulfide dimers can take any value $(0 \le f_{1\mathrm{X}} \le 1)$. Third, since the $C_P$ sulfenic acids quench Trp fluorescence it is assumed that the quantum yield increase associated with the condensation of the sulfenic acid in a 1-disulfide dimer or in an anomalous dimer is no higher than that associated with the condensation of both sulfenic acids $(0 \le \theta_{\mathrm{SO}}, \theta_{\mathrm{SX}} \le 1)$.

With these considerations only one of the six possible assignments yields consistency between theoretical and experimental pre-exponential weights. Namely, $k_{2,\mathrm{SX}} = 7.9 \pm 0.7\ \mathrm{s^{-1}}$, $k_{2,\mathrm{SO}} = 0.94 \pm 0.015\ \mathrm{s^{-1}}$, $k_{2,\mathrm{SS}} = 0.396 \pm 0.004\ \mathrm{s^{-1}}$.

##### 2.2.3 Analysis of the sulfinylation of the C172S Prdx2 mutant

The C172S resolving Cys Prdx2 mutant cannot form interchain disulfides. $H_2O_2$ addition causes an initial fluorescence decrease followed by a gradual recovery that is attributable to sulfinylation (Figure 6B). Analysis of the recovery phase revealed a bi-exponential increase in fluorescence such that the largest characteristic constant is proportional to $H_2O_2$ concentration and the smallest is not. The pre-exponential coefficient corresponding to these characteristic constants were negative and positive, respectively. Below we show that these features are consistent with non-cooperative sulfinylation accompanied by slow bleaching of the sulfinylated sites.

Considering again that all the available Prdx2-$C_P$ thiolates are instantaneously sulfenylated, the simplest system of equations that can describe the dynamics of sulfinylation accompanied by bleaching of the sulfinic acids ($k_{Bl}$) is as follows (Figure S 5A):

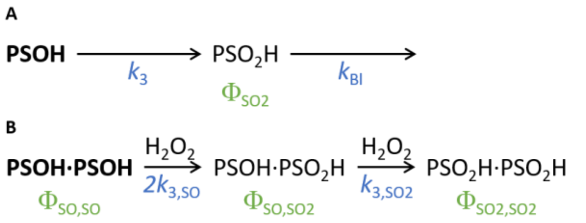

*Figure S 5. Two alternative models to explain the observed Trp fluorescence recovery time course after treatment of 0.5 μM C172S-Prdx2 with supra-stoichiometric $H_2O_2$ concentrations: non-cooperative sulfinylation followed by bleaching (A), and cooperative sulfinylation (B). Species present immediately after initial sulfenylation are indicated in bold. Symbols denoting the fluorescence quantum yields of each species are indicated in green.*

$$\begin{aligned}\frac{d\ PSOH}{d\,t} &= -k_3 PSOH\ H_2O_2 \\ \frac{d\ PSO_2H}{d\,t} &= k_3 PSOH\ H_2O_2 - k_{Bl} PSO_2H\end{aligned} \tag{16}$$

Neglecting the fluorescence quantum yield of the PSOH, the fluorescence signal intensity $\left(I(t)\right)$ will evolve according to:

$$I(t) = B + G\Phi_{SO2}\, PSO_2H(t), \tag{17}$$

where B and G stand for the instrumental baseline and gain (respectively), and $\Phi_{SO2}$ is the fluorescence quantum yield of the $PSO_2H$. Solving equation (16) and replacing into (17) yields:

$$I(t) = B - C\left(e^{-k_3 ct} - e^{-k_{Bl}t}\right), \tag{18}$$

with $C = 2\dfrac{G\Phi_{SO2} Prdx2}{1-\dfrac{k_{Bl}/k_3}{c}}$ and $c = H_2O_{2,0} - 2Prdx2$. Equation (18) constitutes **Model 6**.

Cooperative sulfinylation (Figure S 5B) can also generate a bi-exponential Trp fluorescence recovery. Drawing, where pertinent, on the same assumptions and approximations that underlie equations (10) the corresponding model translates into:

$$\begin{aligned}\frac{d\ PSOH\bullet PSOH}{d\,t} &= -2k_{3,SO} PSOH\bullet PSOH\ H_2O_2 \\ \frac{d\ PSOH\bullet PSO_2H}{d\,t} &= 2k_{3,SO} PSOH\bullet PSOH\ H_2O_2 - k_{3,SO2} PSOH\bullet PSO_2H\ H_2O_2 \\ \frac{d\ PSO_2H\bullet PSO_2H}{d\,t} &= k_{3,SO2} PSOH\bullet PSO_2H\ H_2O_2\end{aligned} \tag{19}$$

with initial conditions $PSOH\bullet PSOH(0) = Prdx2$, $PSOH\bullet PSO_2H(0) = PSO_2H\bullet PSO_2H(0) = 0$.

This model yields:

$$I(t) = B + \Phi_{SO2,SO2} Prdx2 + a_{SO} e^{-2k_{3,SO} c t} + a_{SO2} e^{-k_{3,SO2} c t}, \quad (20)$$

with

$$a_{SO} = -G\ Prdx2 \frac{2k_{3,SO}(\Phi_{SO,SO2} - \Phi_{SO,SO}) - k_{3,SO2}(\Phi_{SO2,SO2} - \Phi_{SO,SO})}{2k_{3,SO} - k_{3,SO2}},$$
$$a_{SO2} = -G\ Prdx2 \frac{2k_{3,SO}}{2k_{3,SO} - k_{3,SO2}}(\Phi_{SO2,SO2} - \Phi_{SO,SO2}). \quad (21)$$

and $\Phi_{x,y}$ the fluorescence quantum yields of the various dimer forms. According to equation (20) -- which defines **Model 7** -- both characteristic constants are $H_2O_{2,0}$-dependent, but either $2k_{3,SO}$ or $k_{3,SO2}$ might be just too small to yield a statistically significant slope of the corresponding characteristic constant *vs. c*. However, algebraic analysis of equations (21) reveals that under these conditions the positive pre-exponential coefficient would be associated with the smallest rate constant (as observed) only if $\Phi_{SO2,SO2} < \Phi_{SO,SO2}$. The latter relationship is very implausible as sulfinylation is associated with a fluorescence *increase* and both sites are sulfinylated under the conditions of the experiment.

Attending to the previous considerations, we fitted Model 6 to the fluorescence recovery time courses for various supra-stoichiometric $H_2O_2$ concentrations, starting at $t = 4$ s, with $B, C, k_B$ and $k_3^* = k_3 c$ as adjustable parameters. These fits yielded a random 0-centered distribution of residuals over the whole time range (Figure 6B). As expected, the best-fit characteristic constants with the positive pre-exponential coefficient, corresponding to $k_{Bl}$, were independent of $H_2O_{2,0}$ (not shown) whereas those with the negative pre-exponential coefficient increased linearly with $H_2O_{2,0}$ (Figure 6C). The slope of the latter dependence yields $k_3 = (3.4 \pm 0.1) \times 10^3$ M$^{-1}$s$^{-1}$. The intercept at the origin (0.0078±0.0007 s$^{-1}$) is statistically significant but

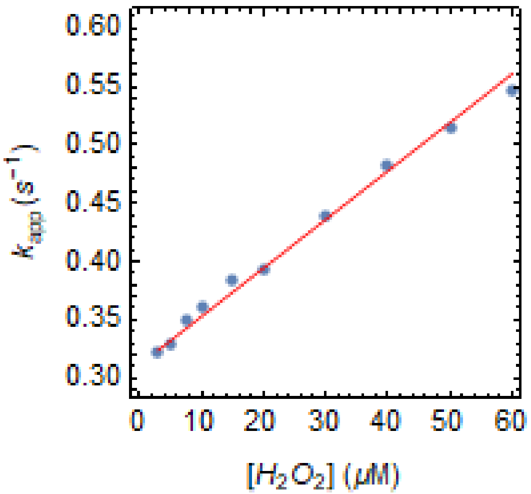

*Figure S 6. Concentration dependence of the characteristic constant for the recovery phase of Trp fluorescence after treatment of Prdx2 (0.5 μM) with excess $H_2O_2$. Slope and intercept at origin were (4.16±0.01)×10$^3$ M$^{-1}$s$^{-1}$ and 0.311±0.002 s$^{-1}$, respectively*

small. Fits of mono-exponential or bi-exponential functions with two independently adjustable pre-exponential coefficients yielded consistently poorer AIC values, demonstrating the superiority of Model 6.

Altogether, these results show that sulfinylation proceeds as a mono-exponential (pseudo-first-order) process. Therefore, sulfinylation at one site in a PSOH·PSOH dimer does not influence the rate constant for sulfinylation at the second site nor the change in fluorescence quantum yield associated to the latter process.

##### 2.2.4 Analysis of the sulfinylation of wild type Prdx2 from Trp fluorescence recovery time courses and gel-based experiments

At high supra-stoichiometric $H_2O_{2,0}$ additions both condensation and sulfinylation contribute to Trp fluorescence recovery and need to be considered in the analysis. Mono-exponential fits of the fluorescence recovery yield characteristic constants that depend linearly on the $H_2O_2$ concentration (Figure S 6).

In order to account for the mutual influences of the state (PSOH, PSS, $PSO_2H$ or PSX) at one site on the condensation and sulfinylation rate constants at the other site we consider the reaction scheme in Figure S 7. As for Model 5 (Section 2.2.2) we will consider here that all the available Prdx2-$C_P$ thiolates are instantaneously sulfenylated upon treatment of Prdx2 with supra-stoichiometric $H_2O_2$. Moreover, for $H_2O_{2,0} >> 2(2-f_1)Prdx2$ the $H_2O_2$ concentration remains approximately constant over the time course of the experiments. Attending to these considerations, the dynamics of sulfinylation is approximated by the following system of ordinary differential equations:

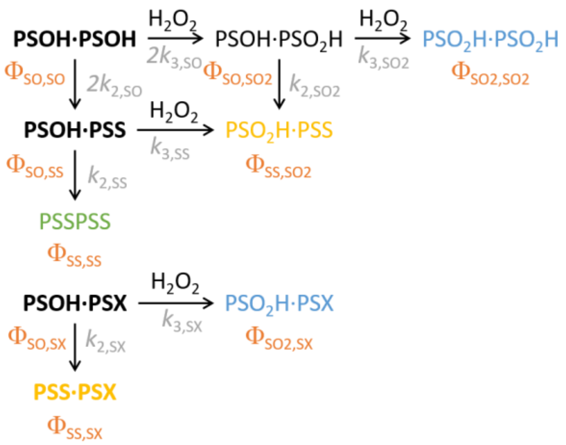

*Figure S 7. Reaction scheme considered for fitting the Trp fluorescence recovery experiments with supra-stoichiometric bolus $H_2O_2$ addition to Prdx2. The species present immediately after sulfenylation are represented in bold type. The species present immediately after sulfenylation are represented in bold type. The final species forming 0-, 1- and 2-disulfide dimers are represented in blue, yellow and green, respectively. The fluorescence quantum yields of the various species and the rate constants are represented in salmon and grey, respectively. This is the same reaction scheme depicted in Figure 6A, repeated here to highlight the quantum yield symbols and number of disulfides.*

$$\begin{aligned}
\frac{d\,PSOH\bullet PSOH}{d\,t} &= -2k_{2,\mathrm{SO}}PSOH\bullet PSOH - 2k_{3,\mathrm{SO}}PSOH\bullet PSOH\;H_2O_2 \\
\frac{d\,PSOH\bullet PSS}{d\,t} &= 2k_{2,\mathrm{SO}}PSOH\bullet PSOH - k_{2,\mathrm{SS}}PSOH\bullet PSS - \\
&\quad - k_{3,\mathrm{SS}}PSOH\bullet PSS\;H_2O_2 \\
\frac{d\,PSS\bullet PSS}{d\,t} &= k_{2,\mathrm{SS}}PSOH\bullet PSS \\
\frac{d\,PSOH\bullet PSO_2H}{d\,t} &= 2k_{3,\mathrm{SO}}PSOH\bullet PSOH\;H_2O_2 - k_{2,\mathrm{SO2}}PSOH\bullet PSO_2H - \\
&\quad - k_{3,\mathrm{SO2}}PSOH\bullet PSO_2H\;\;H_2O_2 \\
\frac{d\,PSO_2H\bullet PSS}{d\,t} &= k_{3,\mathrm{SS}}PSOH\bullet PSS\;H_2O_2 + k_{2,\mathrm{SO2}}PSOH\bullet PSO_2H \\
\frac{d\,PSO_2H\bullet PSO_2H}{d\,t} &= k_{3,\mathrm{SO2}}PSOH\bullet PSO_2H\;\;H_2O_2 \\
\frac{d\,PSOH\bullet PSX}{d\,t} &= -k_{2,\mathrm{SX}}PSOH\bullet PSX\; - k_{3,\mathrm{SX}}PSOH\bullet PSX\;H_2O_2 \\
\frac{d\,PSS\bullet PSX}{d\,t} &= k_{2,\mathrm{SX}}PSOH\bullet PSX \\
\frac{d\,PSO_2H\bullet PSX}{d\,t} &= k_{3,\mathrm{SX}}PSOH\bullet PSX\;H_2O_2
\end{aligned} \tag{22}$$

with the initial conditions $PSOH\bullet PSOH_0 = (1-f_{0X})(1-f_1)Prdx2,$ $PSOH\bullet PSS_0 = (1-f_{1X})\,f_1 Prdx2,$ $PSOH\bullet PSX_0 = f_{0X}\,(1-f_1)Prdx2,$ $PSS\bullet PSX_0 = f_{1X}\,f_1 Prdx2,$ $PSS\bullet PSS_0 = PSO_2H\bullet PSS_0 = PSO_2H\bullet PSO_2H_0 = 0.$

Considering that the time evolution of the fluorescence intensity signal is given by:

$$\begin{aligned} I(t) = B + G(&\Phi_{\text{SO,SO}} PSOH{\bullet}PSOH(t) + \Phi_{\text{SO,SS}} PSOH{\bullet}PSS(t) + \Phi_{\text{SS,SS}} PSS{\bullet}PSS(t) + \\ &+\Phi_{\text{SO,SO2}} PSOH{\bullet}PSO_2H(t) + \Phi_{\text{SS,SO2}} PSO_2H{\bullet}PSS(t) + \Phi_{\text{SO2,SO2}} PSO_2H{\bullet}PSO_2H(t) + \\ &+\Phi_{\text{SO,SX}} PSOH{\bullet}PSX(t) + \Phi_{\text{SS,SX}} PSS{\bullet}PSX(t) + \Phi_{\text{SO2,SX}} PSO_2H{\bullet}PSX(t)), \end{aligned} \tag{23}$$

with $\Phi_{x,y}$ the fluorescence quantum yields of the various dimer forms, and replacing the solution of equations (22) one finds an expression in the form:

$$I(t) = B + C_{\text{SX}} e^{-(k_{2,\text{SX}} + k_{3,\text{SX}} c)t} + C_{\text{SO}} e^{-2(k_{2,\text{SO}} + k_{3,\text{SO}} c)t} + C_{\text{SS}} e^{-(k_{2,\text{SS}} + k_{3,\text{SS}} c)t} + C_{\text{SO2}} e^{-(k_{2,\text{SO2}} + k_{3,\text{SO2}} c)t}. \tag{24}$$

Here, $B$, $C_{\text{SX}}$, $C_{\text{SO}}$, $C_{\text{SS}}$ and $C_{\text{SO2}}$ are relatively complicated functions of the rate constants, quantum yields, initial dimer fractions, and $c$, the excess $H_2O_2$ over Prdx2 active sites. Equation (24) defines **Model 8**.

According to equation (24), in presence of anomalous reduced dimers $(i.e., f_{0\text{X}} > 0)$ a tetra-exponential Trp fluorescence recovery is expected and the rate constants for either condensation or sulfinylation at one site depend on the redox state of the second site in the dimer. At $H_2O_{2,0} > 4Prdx2$ the characteristic constants will increase linearly with $c$, with slopes yielding the sulfinylation rate constants and intercepts at the origin yielding the rate constants for condensation in the various dimer types. Therefore, if less than four significantly different intercepts at the origin are detected, or one of these (corresponding to $2k_{2,\text{SO}}$) is twice another one, this means that some changes in the redox state of one site do not influence the condensation rate constant at the other site. Likewise, if less than four significantly different slopes are detected, or one of these (corresponding to $2k_{3,\text{SO}}$) is twice another one, this means that some changes in the redox state of one site do not influence the sulfinylation rate constant at the other site. Also noteworthy, in the characteristic constants in equation (24) the sulfinylation rate constant for one type of dimer is always associated to the condensation rate constant for the same type of dimer. Consequently, clarifying the correspondence of the latter rate constants to the characteristic constants obtained in an experiment with stoichiometric $H_2O_2$ and otherwise similar conditions will help clarifying the correspondence between the sulfinylation rate constants and the slopes of the characteristic constants with respect to the $H_2O_2$ concentration.

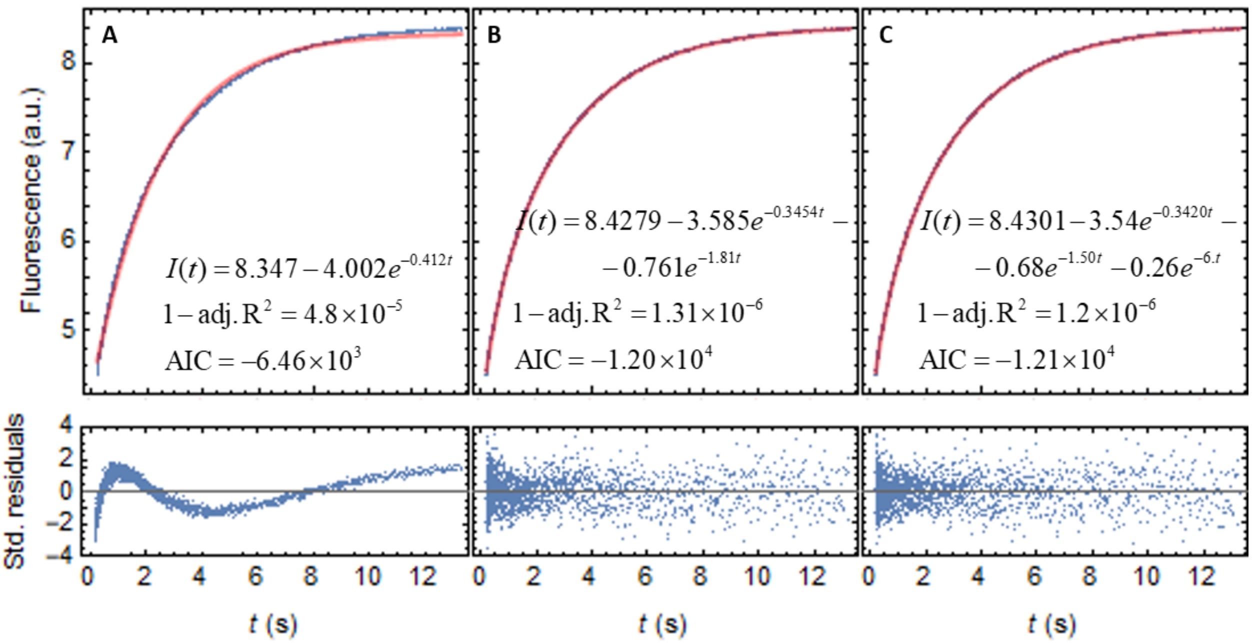

*Figure S 8. Mono- (A), bi- (B) and tri-exponential (C) variance-weighted fits to typical Trp fluorescence recovery time course after treatment of 0.5 µM Prdx2 monomers with 30 µM $H_2O_2$. Red, best-fit curves. Prdx2 was reduced with a five-fold excess of DTT per cysteine the day before experiment. The remaining DTT was removed by filtration (Amicon Ultra 10 kDa, Merk Millipore, Darmstadt, Germany). Protein solutions were stocked in oxygen free atmosphere. $H_2O_2$ was diluted from a stock solution and its concentration was determined spectrophotometrically at 240 nm ($\mathcal{E}_{240nm}$ = 43.6 $M^{-1}.cm^{-1}$). Reactions were performed in 5 mM phosphate buffer pH 7.4 at 25◦C and Prdx2 fluorescence ($\lambda_{ex}$ = 280 nm, emission filter >320 nm) was monitored in a stopped-flow device (Applied Photophysics SX20MV, Leatherhead, United Kingdom).*

With this in mind, we fitted the multi-exponential models to Trp fluorescence recovery time courses for treatment of 0.5 µM Prdx2 monomers with stoichiometric and various supra-stoichiometric $(H_2O_{2,0} > 4Prdx2)$ $H_2O_2$ concentrations according to the workflow described in Figure S 3. This procedure revealed a tri-exponential fluorescence recovery in all cases, with all the pre-exponential coefficients negative (Figure S 8). For stoichiometric $H_2O_2$ addition the best-fit characteristic constants were 3.28±0.04 $s^{-1}$, 1.07±0.03 $s^{-1}$ and 0.2445±0.0004 $s^{-1}$, with pre-exponential weights 0.0703±0.0006, 0.139±0.001 and 0.791±0.001, respectively (means ± standard errors of four replicates). In this experiment the initial fractions of 1-disulfide and anomalous dimers were not determined. For this reason, several alternative assignments of the characteristic constants above to the rate constants in Model 5 permit consistency between experimental and theoretical pre-exponential weights under plausible assumptions $(f_1, f_{0X} \leq 0.2,\ 0 \leq f_{1X} \leq 1,\ 0 \leq \theta_X, \theta_O \leq 1)$. However, only the following assignment is also consistent with the results from the experiment in Section 2.2.2: $k_{2,\mathrm{SX}} = 3.28 \pm 0.04\,\mathrm{s}^{-1}$, $k_{2,\mathrm{SO}} = 0.54 \pm 0.15\,\mathrm{s}^{-1}$, $k_{2,\mathrm{SS}} = 0.2445 \pm 0.0004\,\mathrm{s}^{-1}$.

With supra-stoichiometric $H_2O_2$ additions, all the characteristic constants exhibited the expected linear dependence on $H_2O_{2,0}$ as per equation (24), with slopes *vs.* *c* of $(18.\pm6)\times10^3\ \mathrm{M^{-1}s^{-1}}$, $(7.6\pm1.1)\times10^3\ \mathrm{M^{-1}s^{-1}}$ and $(3.33\pm0.03)\times10^3\ \mathrm{M^{-1}s^{-1}}$, and intercepts at the origin $3.8\pm0.15\ \mathrm{s^{-1}}$, $1.15\pm0.03\ \mathrm{s^{-1}}$ and $0.2435\pm0.0009\ \mathrm{s^{-1}}$, respectively (Figure 6D-F, respectively). The second slope is within experimental error of twice the third, and the corresponding inferred sulfinylation rate constant is remarkably similar to that obtained for the C172S mutant. In turn, the intercepts at the origin are not significantly different from the characteristic constants obtained for stoichiometric $H_2O_{2,0}$ addition. This correspondence supports attributing the slopes above to $k_{3,\mathrm{SX}}$, $2k_{3,\mathrm{SO}}$ and $k_{3,\mathrm{SS}}$, respectively. The fact that no other $H_2O_{2,0}$-dependent characteristic constant is detected indicates that $k_{3,\mathrm{SO2}} \approx k_{3,\mathrm{SO}}$ as inferred from the experiment with C172S Prdx2 as well, and also $k_{2,\mathrm{SO2}} \approx k_{2,\mathrm{SO}}$. These differences among condensation and sulfinylation rate constants translate into significant differences in resistance to sulfinylation $(r_y = k_{2,y} / k_{3,y})$[3] among the various types of sites, by the following order: PSOH·PSX ($r_{\mathrm{SX}}$ = 0.21±0.07 mM), PSOH·PSOH=PSOH·PSO$_2$H ($r_{\mathrm{SO}} = r_{\mathrm{SO2}}$ = 0.15±0.02 mM), PSOH·PSS ($r_{\mathrm{SS}}$ = 73.1±0.7 µM). For the $s_y = k_{3,y} / k_{2,y}$ sulfinylation susceptibilities, which may be more familiar to most readers, the order and values are: PSOH·PSS [$s_{\mathrm{SS}}$ = $(1.37\pm0.01)\times10^4\ \mathrm{M^{-1}}$], PSOH·PSOH=PSOH·PSO$_2$H [$s_{\mathrm{SO}} = s_{\mathrm{SO2}} = (7.\pm1)\times10^3\ \mathrm{M^{-1}}$], PSOH·PSX [$s_{\mathrm{SX}}$ = $(5.\pm2)\times10^3\ \mathrm{M^{-1}}$]. Note that since the results above show that $k_{3,\mathrm{SO2}} \approx k_{3,\mathrm{SO}} \approx k_{3,\mathrm{SS}}$ the difference in resistance to sulfinylation between PSOH·PSOH and PSOH·PSS is essentially a consequence of the cooperativity in condensation.

[3] These ratios represent the $H_2O_2$ concentration at which, for each type of dimer, the sulfinylation rate becomes identical to the condensation rate.

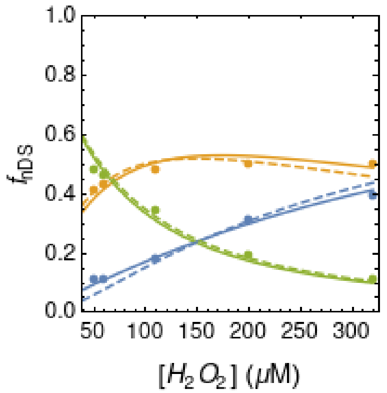

*Figure S 9. Best fit of Model 9 to the relative densities of the monomer (blue), 1-disulfide dimer (yellow) and 2-disulfide dimer (green) bands from non-reducing SDS-PAGE gels for 5 μM Prdx2 treated with supra-stoichiometric $H_2O_2$ concentrations. Solid lines, setting $r_{SO2}=r_{SO}$ and leaving $r_{SO}$, $r_{SS}$ and $r_{SX}$ adjustable; dashed lines, setting $r_{SO2}=r_{SO}=r_{SS}=r_{SX}$ and leaving only the common r adjustable (no cooperativity).*

The cooperativity in condensation can be assessed through an independent experimental method. Namely, by examining how the relative densities of the monomer, 1-disulfide and 2-disulfide bands in a non-reducing SDS-PAGE gel vary as function of supra-stoichiometric $H_2O_2$ concentrations. Because no reduced Prdx remains, the monomer and 1-disulfide bands corresponds to PSO₂H·PSO₂H (dissociated) and PSO₂H·PSS dimers, respectively. Considering the equivalence of $k_{3,SO2}, k_{3,SO}$ and $k_{3,SS}$, the slower condensation of the PSOH·PSS relative to that of the PSOH·PSOH translates into a higher proportion of PSO2H·PSS (1-disulfide) dimers being formed than expected in absence of cooperativity.

The quantitative extent of this effect can be predicted by solving equations (22) analytically to find the final fractions of PSO₂H·PSO₂H ($f_{0,\infty}$), PSO₂H·PSS ($f_{1,\infty}$), and PSS·PSS ($f_{2,\infty}$) dimers, which yields:

$$
\begin{aligned}
f_{0,\infty} &= (1-f_1)\left( (1-f_{0X})\frac{\frac{h}{g_{SO}r_{SS}}}{1+\frac{h}{g_{SO}r_{SS}}}\frac{\frac{h}{g_{SO2}r_{SS}}}{1+\frac{h}{g_{SO2}r_{SS}}} + f_{0X}\frac{\frac{h}{r_{SX}}}{1+\frac{h}{r_{SX}}} \right) \\
f_{1,\infty} &= 1 - f_{0,\infty} - f_{2,\infty} \\
f_{2,\infty} &= \left( (1-f_{1X})f_1 + (1-f_{0X})(1-f_1)\frac{1}{1+\frac{h}{g_{SO}r_{SS}}} \right)\frac{1}{1+\frac{h}{r_{SS}}}
\end{aligned}
\qquad (25)
$$

with $h = H_2O_{2,0} - (2 - f_{0X}(1-f_1) - (1+f_{1X})f_1)Prdx2$, $g_{SO} = r_{SO}/r_{SS} = k_{2,SO}/k_{2,SS}$ and $g_{SO2} = r_{SO2}/r_{SS} = k_{2,SO2}/k_{2,SS}$. Equations (25) define **Model 9**.

We determined the initial fraction of 1-disulfide $(f_1 = 0.06)$ and anomalous $(f_X = f_{0X}(1-f_1) + f_{1X}f_1 = 0.12)$ dimers from the relative densities of the 1-disulfide dimer bands of the gel lanes corresponding to 0 and equi-stoichiometric $H_2O_2$ additions to 5 μM total

Prdx2, respectively. We have also assumed that the fraction of anomalous 0-disulfide and 1-disulfide dimers is identical ($f_{0X} = f_{1X} = 0.12$), and we set $r_{SO2} = r_{SO}$, as indicated by the Trp fluorescence recovery experiments. We thus fitted the data with the three adjustable parameters $g_{SO}$, $r_{SS}$ and $r_{SX}$. This procedure yielded a very good fit (Figure S 9, solid lines) and the following best-fit estimates: $g_{SO} = 1.8 \pm 0.2$, $r_{SS} = (1.14 \pm 0.08) \times 10^2$ µM, $r_{SX} = 32. \pm 16.$ µM (Adj. $R^2$= 0.996, AICc=-62.). This estimate for $g_{SO}$ is not only significantly higher than 1, but also not significantly different from the $g_{SO} = k_{2,SO} / k_{2,SS}$ ratio obtained from the Trp fluorescence recovery experiment, reinforcing the evidence for negative cooperativity in condensation. Moreover, a model assuming non-cooperative condensation ($g_{SO} = 1$) yields a best fit ($r_{SS} = (1.53 \pm 0.07) \times 10^2$ µM, $r_{SX} = (3. \pm 1.) \times 10^2$ µM, adj. $R^2$= 0.993, AICc= 56. Figure S 9, dashed lines) with a significantly lower likelihood ($p<0.045$, based on the difference in AICc), which underestimates the preference for forming PSO2H·PSS dimers *vs*. di-sulfinylated dimers at high $H_2O_2$ concentrations. Distinct assumptions about the distribution of anomalous dimers by the 0- and 1-disulfide fractions do not alter the conclusions above.

##### 2.2.5 A note on alternative explanations for the multi-exponential fluorescence recovery

Any alternative model to explain the Trp fluorescence recovery data presented in this paper must be consistent with all the following observations:

1. The fluorescence recovery associated with the condensation of the sulfenic acids is tri-exponential, with all the pre-exponential coefficients negative.
2. The fluorescence recovery associated with the hyperoxidation of WT Prdx2 is also tri-exponential, the slopes of the dependence of the characteristic constants on the $H_2O_2$ concentration are distinct, and the values of the intercept at the origin match the values of the characteristic constants associated to condensation.
3. The slope of the $H_2O_2$ concentration dependence of the second slowest characteristic constant is about twice that of the slowest characteristic constant.

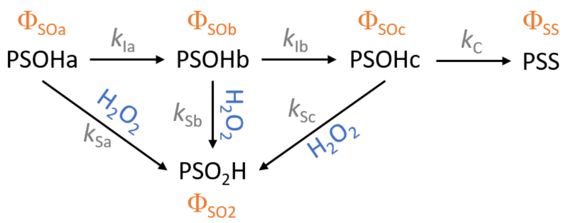

*Figure S 10. Alternative model to explain the fluorescence recovery results. Fluorescence quantum yields of the various species and rate constants are indicated in salmon and gray, respectively.*

4. All the pre-exponential coefficients of the fluorescence recovery associated to hyperoxidation are negative.
5. The fluorescence recovery associated to sulfinylation of the C172S Prdx2 mutant is mono-exponential (after removing the component due to bleaching), and the characteristic constant increases linearly with the concentration of added $H_2O_2$.
6. The slope of this $H_2O_2$ concentration dependence is identical to that for the slowest characteristic constant for the fluorescence recovery associated to the hyperoxidation of WT Prx2.

An independent-sites model such that condensation can only occur after a series of slow conformational changes of the sulfenic form but all of the intermediate conformations can be sulfinylated (Figure S 10) is consistent with observations 1-4 for some choices of parameter values. Namely, the following two rate constant assignments to the slopes and intercepts at the origin for the $H_2O_2$ concentration dependence of the characteristic constants for the fluorescence recovery of the WT Prx2 exposed to excess $H_2O_2$ are consistent with the set of pre-exponential weights for plausible values of $\theta_{ba} = \frac{\Phi_{SS} - \Phi_{SOb}}{\Phi_{SS} - \Phi_{SOa}}, \theta_{ca} = \frac{\Phi_{SS} - \Phi_{SOc}}{\Phi_{SS} - \Phi_{SOa}}$ and $\delta = \frac{\Phi_{SO2} - \Phi_{SS}}{\Phi_{SS} - \Phi_{SOa}}$: $k_{Sa}$ = (18.±6)×10$^3$ M$^{-1}$s$^{-1}$, $k_{Sb}$ = (7.6±1.1)×10$^3$ M$^{-1}$s$^{-1}$, $k_{Sc}$ = (3.33±0.03)×10$^3$ M$^{-1}$s$^{-1}$, $k_{Ia}$ = 3.8±0.15 s$^{-1}$, $k_{Ib}$ = 1.15±0.03 s$^{-1}$ and $k_C$ = 0.2435±0.0009 s$^{-1}$, or $k_{Sa}$ = (7.6±1.1)×10$^3$ M$^{-1}$s$^{-1}$, $k_{Sb}$ = (18.±6)×10$^3$ M$^{-1}$s$^{-1}$, $k_{Sc}$ = (3.33±0.03)×10$^3$ M$^{-1}$s$^{-1}$, $k_{Ia}$ =1.15±0.03 s$^{-1}$, $k_{Ib}$ = 3.8±0.15 s$^{-1}$ and $k_C$ = 0.2435±0.0009 s$^{-1}$.

However, it is difficult to explain observations 5 and 6 in this framework. Because the slope of the $H_2O_2$ concentration dependence of the characteristic constant for the fluorescence recovery of the C172S mutant is identical to $k_{Sc}$, one ought to explain why would the C172S mutation render the Ia, Ib, Sa and Sb steps undetectable. Moreover, whereas Model 8 offers a natural explanation for observations 3 and 6, this alternative model does not. Finally, no independent-sites model can explain the preference for forming PSO2H·PSS dimers highlighted in the SDS-PAGE experiment described in the previous section.

###### 2.2.6 Analysis of the pH dependence of the characteristic constants for fluorescence recovery

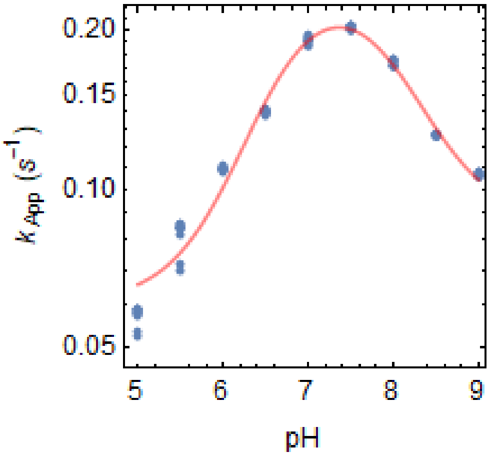

*Figure S 11. pH dependence of the characteristic constant obtained from mono-exponential fits to the Trp fluorescence recovery after treatment of 1 µM Prdx2 with 1 µM or 2 µM $H_2O_2$ (representative experiment). The dots represent best-fit estimates of the characteristic constants for replicate injections from two experiments. The solid line represents the best-fit curve for the pH-dependence model described in ref. (12).*

Another fundamental distinction between the cooperativity and the serial isomerization models above is that only in the former model the characteristic constants for fluorescence recovery reflect processes that are tightly coupled to the condensation reaction proper. Therefore, only the former model predicts that all the characteristic constants respond similarly to factors that are expected to influence the rate constant for condensation. One such factor is pH. Based on mono-exponential fits to the Trp fluorescence recovery after treatment of Prdx2 with excess $H_2O_2$, Portillo-Ledesma *et al.* (12) have shown that the characteristic constant shows a bell-shaped pH dependence. This dependence is well fitted by a model that considers the contributions of the reactions

$$C_PSOH + C_RS^- \underset{k_{2A}}{\rightarrow} C_PSSC_R + OH^- \tag{26}$$

$$C_PSO^- + C_RS^- \underset{k_{2B}}{\rightarrow} C_PSSC_R + OH^- \tag{27}$$

$$C_PSOH + C_RSH \underset{k_{2C}}{\rightarrow} C_PSSC_R + H_2O \tag{28}$$

for the observed resolution rate, accounting for the $pK_a$ of the sulfenic acid and the resolving thiol. We therefore determined the pH dependence of the characteristic constants obtained from tri-exponential fits to the fluorescence recovery. In our experiments, the mono-exponential fits yielded similar results to ref. (12) (Figure S 11 and Table S 2). And, remarkably, all the three characteristic constants obtained from tri-exponential fits showed a similar bell-shaped pH depen-

*Table S 2. Estimates of parameters of the pH-dependence model obtained from mono-exponential or tri-exponential fits to the Trp fluorescence recovery after treatment of 1 µM Prdx2 with 1 µM or 2 µM $H_2O_2$.*

| **Parameter** | **Mono-exponential** | **Tri-exponential** | | |
|---|---|---|---|---|
| | | **Slowest** ($k_{2.,SS}$) | **Intermediate** ($2k_{2.,SO}$) | **Fastest** ($k_{2.,SX}$) |
| $k_{2A}$ ($s^{-1}$) | 8.±1. | 3.1±0.5 | 14.±8. | (1.±0.9)×$10^2$ |
| $k_{2B}$ ($s^{-1}$) | 0.087±0.003 | 0.063±0.002 | 0.42±0.03 | 1.1±0.2 |
| $k_{2C}$ ($s^{-1}$) | 0.061±0.002 | 0.018±0.003 | 0.20±0.03 | 0.6±0.2 |
| $pK_a(C_P\text{-SOH})$ | 6.57±0.04 | 6.82±0.04 | 7.0±0.1 | 6.8±0.2 |
| $pK_a(C_R\text{-SH})$ | 8.06±0.05 | 7.88±0.04 | 8.1±0.2 | 8.2±0.3 |
| Adj. $R^2$ | 0.998 | 0.9991 | 0.994 | 0.98 |

dence that is well fitted by the pH-dependence model from ref. (12) (Figure 6A). (The underlying assumption for applying this model separately to each characteristic constant is that de/protonation of the $C_P$-SOH and $C_R$-SH equilibrates rapidly, compared to the time scale of resolution.) The best-fit parameter estimates for a representative experiment are shown in Table S 2. Importantly, the $pK_a$s of the sulfenic acid and of the resolving thiol estimated from the pH dependence of the three characteristic constants are not significantly different, as expected from similar processes that are tightly coupled to the condensation reaction. In turn, although the condensation of a sulfenic acid in a PSOH·PSOH dimer does not significantly influence the $pK_a$s, it does substantially decrease the rate constants for reactions (26)-(28).

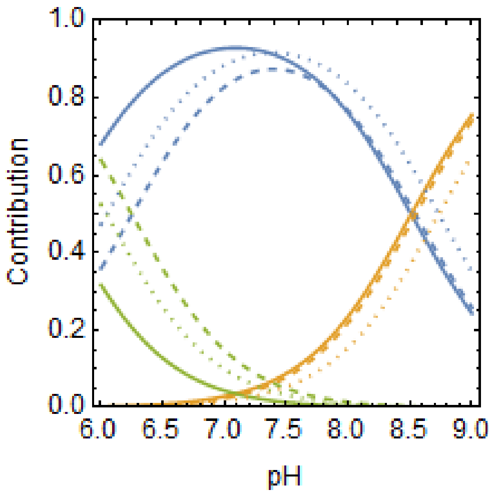

*Figure S 12. pH dependence of the relative contributions of reactions (26) (blue), (27) (yellow) and (28) (green) for the rate of condensation in PSOH·PSS (solid lines), PSOH·PSOH (dashed lines) and PSOH·PSX (dotted lines) dimers.*

These results also allow estimating the relative contributions of reactions (26)-(28) for the overall rate of condensation in each type of dimer (Figure S 12) and the pH dependence of cooperativity (Figure 6B). At pH above 7.5 the relative contributions of the three reactions for the condensation rate are similar between PSOH·PSOH and PSOH·PSS dimers. However, below pH 7.5 reaction (26) contributes substantially more, and reaction (28) substantially less, for the condensation rate of PSOH·PSOH dimers than for that of PSOH·PSS dimers. Cooperativity is weakest around pH 7.0 and stronger away from neutral pH.
